# CellPace: A temporal diffusion-forcing framework for simulation, interpolation and forecasting of single-cell dynamics

**DOI:** 10.64898/2026.02.25.707938

**Authors:** Chen Su, Amin Emad

## Abstract

Single-cell omics technologies resolve cellular heterogeneity at high resolution but provide only static snapshots of continuous developmental processes. This makes it difficult to recover coherent temporal dynamics when developmental stages are irregularly sampled or missing. While recent generative models can simulate observed cell states, they often treat timepoints as discrete categories, hindering interpolation across gaps and extrapolation to unobserved future stages. We present CellPace, a generative model that learns and generates developmental dynamics by leveraging a transformer-based temporal diffusion backbone conditioned on continuous, gap-aware temporal encodings. Across diverse mouse developmental lineages, CellPace achieves state-of-the-art performance in simulation, interpolation, and forecasting tasks. Beyond recovering global population statistics, generated cells preserve fine-grained biological structure, retaining dynamic gene regulatory programs and mapping accurately to anatomical regions in spatial transcriptomics data. Furthermore, CellPace extends naturally to multi-modal data, modeling joint RNA-chromatin dynamics even when temporal ordering is inferred from pseudotime. Together, these results position CellPace as a robust framework for modeling and generating continuous developmental dynamics from sparse, cross-sectional single-cell data.

## Introduction

Single-cell sequencing technologies have transformed our view of biological systems by resolving cellular heterogeneity at an unprecedented resolution. However, these assays are inherently destructive. Each cell is measured only once, providing a static snapshot rather than a continuous record of its molecular history. In most developmental studies, cells are sampled at discrete, often irregularly spaced timepoints [1]. Each timepoint is represented by an independent population rather than individual cells followed over time. Intermediate states may be sparsely sampled or entirely absent, and experimental constraints frequently limit the number of stages that can be profiled. As a result, the underlying process (a continuous, stochastic progression of cell states) is only indirectly observed through a set of disjointed, cross-sectional measurements. Reconstructing a coherent temporal dynamics from these independent snapshots remains a central challenge in computational single-cell biology [2].

A range of computational frameworks have been proposed to address this problem. Trajectory inference methods such as PAGA [3] and CellRank2 [4] excel at ordering observed cells along latent progressions and estimating lineage probabilities. Optimal Transport (OT) based frameworks provide a way to align population-level distributions across conditions [5]. These approaches are widely used to describe developmental landscapes and reconstruct lineage relationships. However, they are mainly descriptive: they organize existing cells and infer transitions but typically do not generate molecular profiles for missing states or predict populations beyond the measured timepoints. Models based on continuous-time neural differential equations such as scNODE [6] and scIMF [7] go one step further by learning vector fields that can be integrated forward in time. Yet they still require real cells at a starting timepoint instead of sampling from noise. Furthermore, they usually assume autonomous dynamics in which each cell evolves independently. This assumption may miss population-level interactions and the effects of multiple conditions. More recently, deep generative models such as scDiffusion [8], cfDiffusion [9] and CFGen [5] have been proposed to generate cells from noise. However, these frameworks fundamentally treat time as a categorical label. Designed originally for class-conditional generation, they view developmental stages as discrete, independent conditions rather than points along a continuous axis. Because they do not encode the temporal distance or relationship between stages, they lack the intrinsic logic required to interpolate missing intermediates or extrapolate to future timepoints. Consequently, while they can reproduce cells at observed stages, they cannot model the continuous temporal dynamics that connect them.

Here, we introduce CellPace, a transformer-based diffusion framework for learning continuous developmental dynamics from single-cell data and generating new datapoints pertaining to seen or unseen timepoints. CellPace utilizes temporal diffusion forcing (TDiF), a novel transformer-based diffusion architecture we designed to relax the requirement for regularly sampled time points [10] and enable effective application in single-cell molecular biology. By combining a stable latent representation with a causally masked temporal diffusion model, conditioned on continuous, gap-aware temporal encodings, CellPace can be trained on irregularly sampled snapshots and learn population-level dynamics across arbitrary temporal intervals. This enables it to generate data at observed and unobserved (intermediate and future) timepoints.

We evaluate CellPace across multiple single-cell transcriptomic developmental datasets, demonstrating that it can simulate realistic data at observed timepoints and, more importantly, interpolate intermediate stages and extrapolate to previously unseen future timepoints. We show that CellPace generalizes across time, preserves biological structure, and maintains biologically meaningful gene regulatory dynamics. It accurately preserves positional identity, marker expression, spatial localization, and global topological connectivity, even when intermediate stages are missing. Finally, we show that CellPace can be extended to paired RNA–ATAC data, successfully modeling continuous multiomic developmental dynamics, and can leverage pseudotime to analyze snapshot datasets lacking explicit temporal measurements. To our knowledge, CellPace is the first diffusion-based framework that simultaneously supports simulation from noise, interpolation, and temporal extrapolation for single-cell transcriptomic and multiomic data.

## Results

### CellPace is a generative framework for modeling continuous single-cell temporal dynamics

CellPace is a deep generative model that learns continuous single-cell dynamics from discretely sampled transcriptomic snapshots, enabling simulation, interpolation, and temporal extrapolation (Fig. 1a-1d). During training, it receives transcriptomic profiles of cells at known developmental stages and encodes them into a low-dimensional latent space using a pretrained variational autoencoder (VAE) [11]. To ensure the model does not rely on the assumption of uniformly-spaced intervals, latent embeddings are randomly sampled into training sequences with varying time gaps. To represent this irregular temporal structure explicitly, each position in the sequence is annotated with a two-dimensional vector consisting of the (normalized) temporal position and the elapsed time relative to the previous stage. These gap-aware continuous features enable the model to capture both short-range transitions and long-range developmental dynamics.

**Fig. 1.**
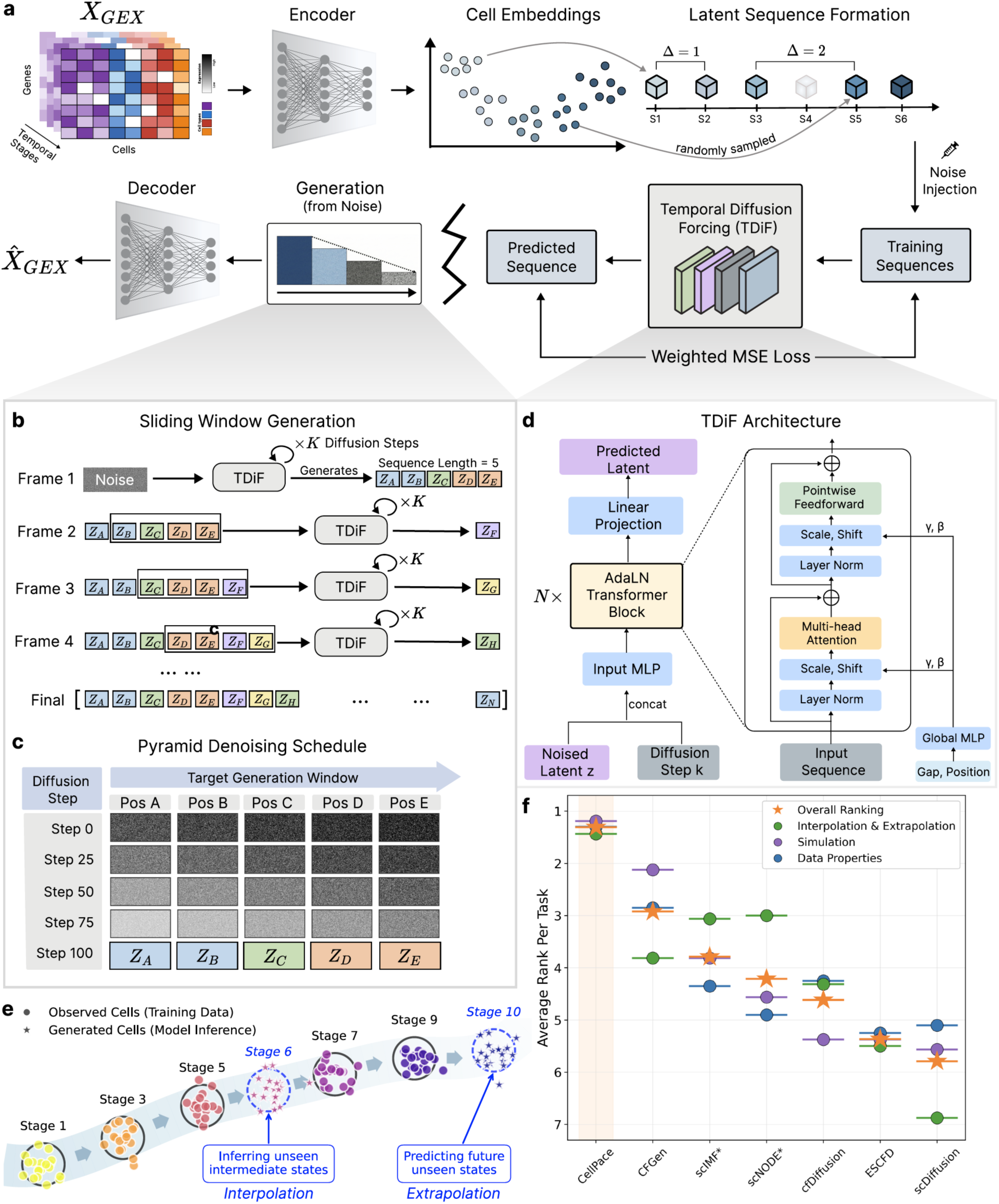
Overview of the CellPace framework. **a,** Schematic representation of the two-stage generative pipeline. Input gene expression (*X_GEX_*) is encoded into cell embeddings via a pretrained variational autoencoder (scVI or MultiVI). Latent sequences are formed by sampling cell embeddings across temporal stages with variable gaps (Δ). A diffusion-based model (Temporal Diffusion Forcing, TDiF) is trained to denoise corrupted latent sequences, enabling iterative generation from pure Gaussian noise at inference time. **b,** Sliding-window strategy enabling arbitrary-length sequence generation, where a fixed-length latent context (e.g., *Z*_*A*_, …, *Z*_*E*_) is shifted forward autoregressively to generate subsequent latent states (e.g., *Z*_*F*_) using previously generated latents as conditioning context. **c,** Pyramid denoising schedule designed for causal temporal modeling via cascaded noise removal, in which earlier sequence positions are resolved prior to later ones to condition future predictions on refined estimates of past states. **d,** TDiF network architecture used during both training and inference, where a global MLP processes biological context, specifically normalized stage positions and inter-stage time gaps, to generate scale (*γ*) and shift (*β*) parameters that modulate transformer blocks and condition denoising on developmental time. **e,** Temporal generalization tasks, illustrating CellPace’s ability to infer valid cell states beyond the training distribution. Interpolation reflects the reconstruction of missing intermediate stages (e.g., Stage 6), while extrapolation (i.e., temporal forecasting) reflects the prediction of future unseen states (e.g., Stage 10) by propagating learned dynamics forward in time. **f,** Benchmarking summary, comparing CellPace with six baseline methods across four developmental datasets. Methods are ranked (1 = best) across three evaluation categories: Interpolation & Extrapolation on held-out stages (green), Simulation on training stages (purple), and Data Properties assessing statistical fidelity (blue). The Overall Ranking (orange star) denotes the average rank across all categories. See Supplementary Tables 1 and 2 for the value of the metrics.

Each sequence is then fed into a module that we call Temporal Diffusion Forcing (TDiF). TDiF builds on the Diffusion Forcing architecture [10] by introducing capabilities, such as handling irregular time intervals, that are required for modeling and generating temporal single-cell transcriptomic data (Fig. 1d). In this module, noise is added to the latent vectors independently for each position according to a cosine variance schedule. This creates varying information density across the sequence: heavily corrupted frames serve as prediction targets, while nearly clean frames provide reliable context. To process this input, the diffusion module employs a Transformer backbone with causal attention masks that prevent information leakage from future states, strictly enforcing the temporal directionality. Meanwhile, the gap-aware temporal features condition the model via Adaptive Layer Normalization (AdaLN) [12], modulating neural activations at each layer according to both developmental position and inter-stage spacing to account for irregular sampling intervals (Fig. 1c). The model is trained to reconstruct clean latent representations from noisy inputs using a mean squared error objective, with Min-SNR weighting [13] to balance optimization across noise levels.

At inference time, CellPace generates latent sequences from noise, conditioned on specified developmental stages. To handle sequences that exceed the fixed training window length, a sliding window approach is utilized to generate the full sequence iteratively, shifting the context forward step-by-step. Within each window generation, instead of denoising all timepoints synchronously, the model employs a Pyramid Schedule [10] (Fig. 1b-1c) in conjunction with a Denoising Diffusion Implicit Models (DDIM) sampler [14] for accelerated inference. The pyramid schedule allocates denoising effort asymmetrically across timepoints, stabilizing earlier stages first so that the reconstructed past provides clean context for generating future states. In each sampling step, TDiF estimates the original clean data; the DDIM update rule then combines this prediction with a derived noise term to compute the next latent state. Once the latent sequence is fully generated, the frozen VAE decoder maps the latent vectors back to high-dimensional gene expression counts (Fig. 1a).

Benchmarking (using different datasets and metrics) against various alternative models, ranging from diffusion and flow matching approaches to neural ordinary and stochastic differential equations (ODE/SDE), show that CellPace achieves superior performance in generating realistic samples from noise in simulation (generating samples similar to the training set), in interpolation (generating samples corresponding to unseen intermediate timepoints), and in extrapolation (predicting future cellular states beyond the range of the training data) (Fig. 1e-1f). Fig. 1f and Supplementary Fig. 1 provide the summary ranking of CellPace compared to other methods on the above three tasks that will be discussed in detail, and using additional analyses, in subsections below.

While CellPace was primarily designed for single-cell transcriptomic simulation, interpolation, and extrapolation across known developmental time, we show that it can be adapted to support multi-omics data (including 10x Genomics RNA–ATAC multiome measurements) and generation from pseudotime.

### CellPace accurately simulates retinal progenitor populations at observed developmental stages

Since CellPace is a generative model, it can generate new transcriptomic profiles from noise. We first asked whether the generated profiles, corresponding to the developmental stages that it has seen during training, capture the distribution and temporal structure of real experimental data (a requirement for its utility as a simulator). We focused on data corresponding to the temporal progression of mouse retinal progenitor cells (RPC) from a recent study [15]. RPCs are multipotent cells that divide and differentiate into multiple retinal cell types, including photoreceptors (rods and cones), horizontal cells, amacrine cells and ganglion cells, as the layered retina is assembled during development [16, 17]. We used the somite count as the temporal measure of the developmental stage and retained 3,000 highly variable genes (HVG) for analysis. The RPC dataset contained 35,275 cells from 31 stages (with somite counts ranging between 0 to 34). We used 28,389 cells from 26 stages for training and validation, while five stages were completely held out as the test set to assess interpolation and extrapolation (discussed in subsequent sections).

We generated cells using CellPace from noise, by conditioning on the target somite index and the inter-stage temporal gap. Fig. 2a visually shows that CellPace generates realistic cells in different developmental stages. Quantitative evaluation further confirmed this observation: comparing real and generated cells using different metrics, CellPace demonstrated the strongest overall performance relative to six state-of-the-art models, comprising latent diffusion-based methods [8, 9, 18], a flow-matching model [5], and continuous-time neural ODE/SDE methods [6, 7] (Fig. 2b-2c). CFGen [5], a recent flow-matching model, also performed well in terms of realistic simulation, compared to other baselines. We repeated the analyses above on three other datasets pertaining to the development of epithelial cells, endothelial cells, and posterior embryo, which consistently showed a high degree of similarity between CellPace-generated and real cells across different stages (Supplementary Fig. 2).

**Fig. 2.**
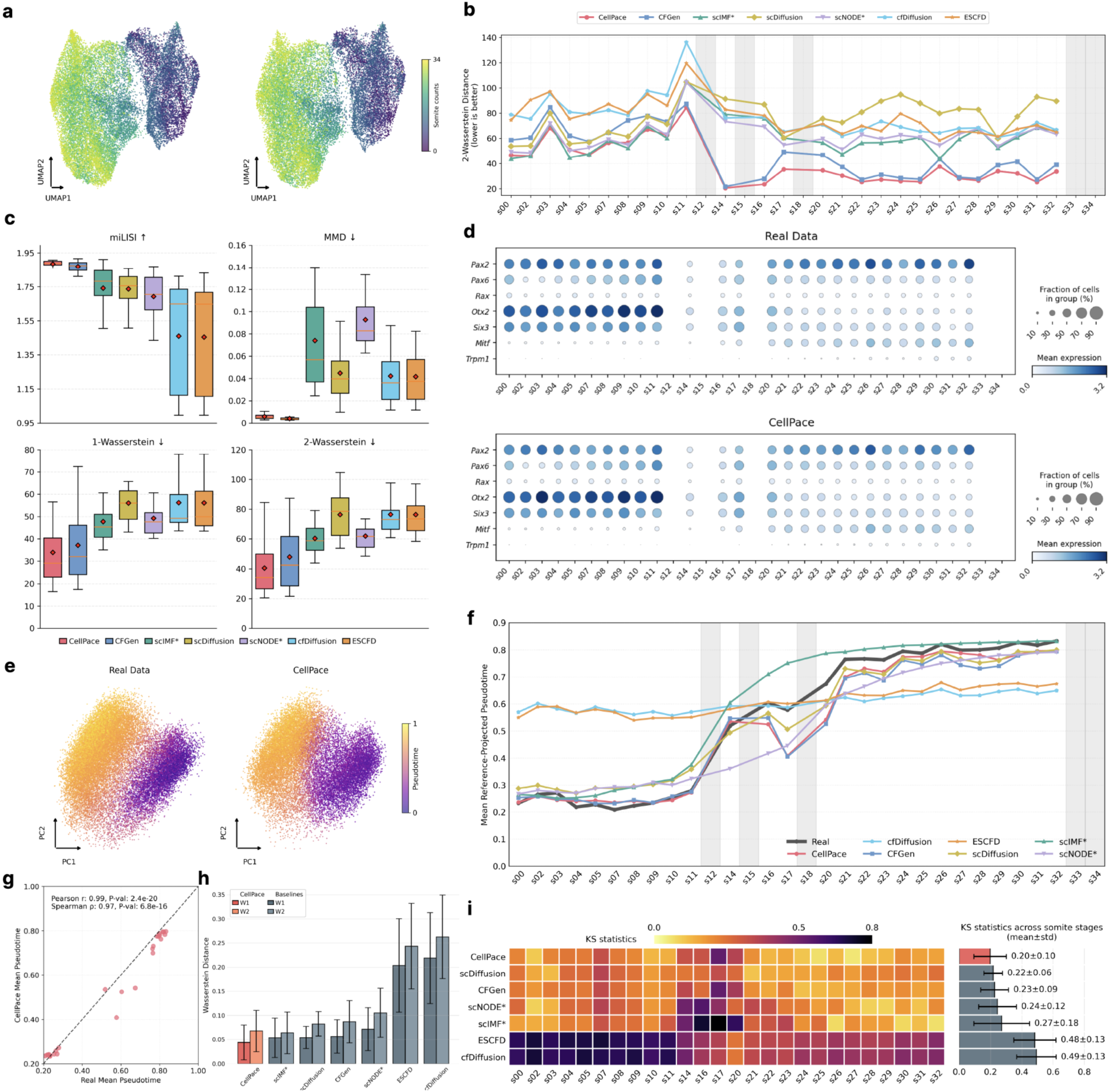
Simulation performance on training stages in retinal progenitor cell (RPC) data. **a,** Joint UMAP embedding of real and CellPace-generated RPC profiles across training stages, colored by somite stage, illustrating global structure and continuity in the latent space. **b,** Temporal consistency analysis quantified by the 2-Wasserstein distance across developmental stages, determined based on somite counts. Grey shaded regions denote held-out stages used for interpolation and extrapolation tasks in subsequent sections. **c,** Distribution of quantitative performance metrics across training somite stages. Box plots summarize per-somite values for miLISI, MMD, 1-Wasserstein, and 2-Wasserstein distances for CellPace and baseline methods. **d,** Marker gene expression dynamics across retinal development. Dot plots compare mean expression (color intensity) and fraction of expressing cells (dot size) between real data (top) and CellPace-generated cells (bottom). Genes are grouped by functional role: stable RPC identity markers (*Pax2*, *Pax6*, *Rax*); early eye-field regulators (*Otx2*, *Six3*) exhibiting decreasing expression; and late retinal pigment epithelium (RPE) markers (*Mitf, Trpm1*) exhibiting increasing expression across stages. Color scales are normalized to the maximum expression observed in the real data for each gene. **e,** Principal component analysis (PCA) projection of real (left) and CellPace-generated (right) cells. Cells are colored by diffusion pseudotime, computed on real data and transferred to generated cells by k-nearest neighbor (k-NN) regression in the shared PCA space. **f,** Mean reference-projected pseudotime across training stages. The solid black line represents the real data trajectory; colored lines represent projected pseudotime using generated data from CellPace and baseline methods. Grey shaded regions indicate held-out test somite stages. **g,** Scatter plot comparing mean pseudotime per somite between real data (x-axis) and CellPace (y-axis). The dashed line indicates identity (y=x). Pearson (r) and Spearman (ρ) correlation coefficients and p-values are reported. **h,** Bar plot of mean 1-Wasserstein (W1) and 2-Wasserstein (W2) distances between real and generated pseudotime distributions for each method, averaged across training somite stages. Error bars indicate standard deviation. **i,** Kolmogorov-Smirnov (KS) statistics comparing real and generated pseudotime distributions. The heatmap (left) shows KS values across individual somite stages for each method, while the bar plot (right) summarizes the mean and standard deviation across stages. Note that * shows methods that instead of generating from noise, required the real data in the initial stage for simulation.

To validate the model beyond global distributions, we inspected gene-expression dynamics for a small panel of RPC canonical markers. The patterns observed in data generated by CellPace closely matched the real data (Fig. 2d). RPC identity genes such as *Rax* and *Pax2* remained broadly expressed across all somite stages; *Otx2* and *Six3* decreased over time, while *Mitf* and *Trpm1* showed the expected late increase associated with retinal pigment epithelium differentiation. The close agreement in both mean expression and fraction of expressing cells indicates that CellPace closely mirrors the underlying developmental progression in the marker-gene space.

Next, we set out to assess how well CellPace preserves the temporal dynamics of RPC development. Fig. 2e visually shows a high concordance between the diffusion pseudotime (DPT) [19] of real and CellPace-generated cells. We used a reference-based trajectory analysis to systematically evaluate this observation. We computed the pseudotime of real training data and transferred it to generated cells via k-nearest-neighbour projection in a shared principal component analysis (PCA) space (Methods). Fig. 2f shows the mean pseudotime as a function of the somite count for the real dataset, as well as data generated by all models. CellPace aligned well with the real data (Pearson r = 0.98, p-value = 2.4E-20) and achieved the lowest 1-Wasserstein (W1) and second lowest 2-Wasserstein (W2) distance (after scIMF). However, it is important to note that unlike CellPace whose data was generated from noise, scIMF required (and received) real cells from the initial timepoint as the starting point for generation. To assess local fidelity, we computed the Kolmogorov-Smirnov (KS) statistics comparing pseudotime distributions within each somite (Fig. 2i), which revealed CellPace to have the smallest mean across all models. Collectively, these analyses show that CellPace accurately captures the temporal structure of the real data.

### CellPace exhibits strong performance in temporal interpolation and forecasting

We next asked whether CellPace could generate transcriptomic profile of cells for intermediate unseen states (interpolation) as well as temporal forecasting (extrapolation). Due to its architecture and design, CellPace learns the temporal relationship of samples using an ordinal or continuous variable of time (e.g., somite number to represent developmental stage), and supports generation in unseen timepoints. Among the models discussed earlier, scNODE and scIMF support generation in arbitrary timepoints via ODE/SDE integration but require initial real cells as input. scDiffusion is the only other model that natively supports interpolation but does not support extrapolation for temporal forecasting. CFGen, ESCFD, and cfDiffusion are designed to be trained with categorical variables and do not support interpolation or extrapolation.

To enable comparison against models that do not allow predictions for unseen stages, we utilized a post-hoc approach: we obtained intermediate-stage predictions by pairwise interpolation between neighbouring training stages, and we generated future-stage predictions by autoregressive linear extrapolation from the two most recent observed stages. For the extrapolation task, we used the last two stages (somite count of 33 and 34) as the test set. For interpolation, however, we carefully analyzed different stages of the RPC dataset and selected three stages that reflected different levels of prediction difficulty as our test sets (Methods): easy (somite count of 18), moderate (somite count of 15), and hard (somite count of 12).

Visually, CellPace-generated cells densely populated the retinal progenitor manifold at all three held-out interpolation stages as well as the two extrapolation stages: red (generated) circles are closely interleaved with the grey reference cells, preserving the overall shape and density of the UMAP embedding (Fig. 3a). By contrast, other methods, even CFGen that performed well in simulation, struggled in generating realistic data (both visually and quantitively), particularly in the moderate (S15) and hard (S12) interpolation stages and in the extrapolation stages (S33 and S34) (Fig. 3a-3b). CellPace showed a strong extrapolation performance, achieving the lowest W2 distance among all methods and a consistent high miLISI score. Interestingly, scIMF also achieved a strong extrapolation miLISI score, yet it had a substantial higher W2 distance compared to CellPace in both extrapolation and interpolation. We repeated the analyses above using three other datasets, which resulted in similar patterns (Supplementary Fig. 3): CellPace-generated data showed the lowest W2 distance with real data among all models, and CellPace and scIMF consistently outperformed other baselines in terms of miLISI.

**Fig. 3.**
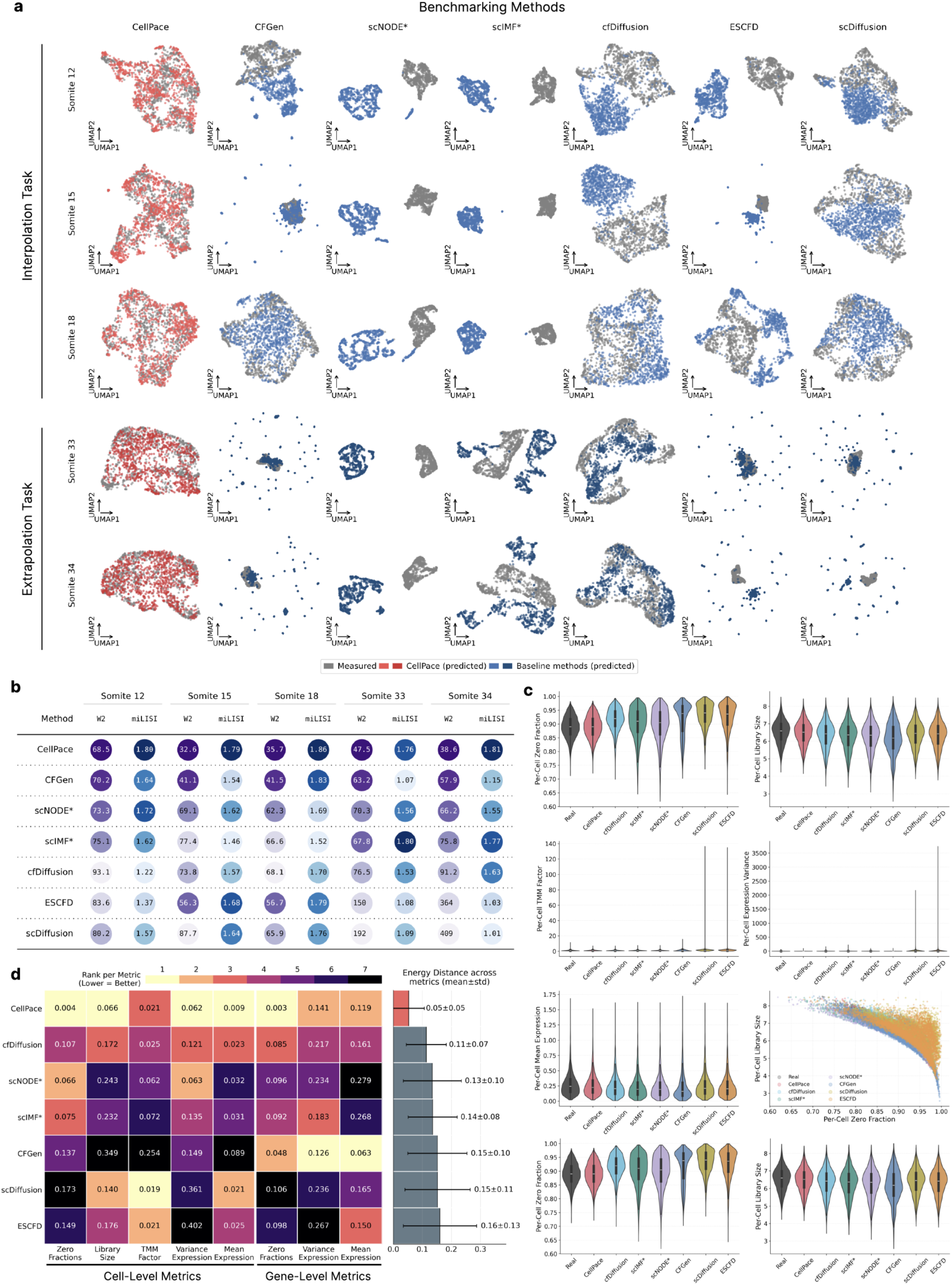
Interpolation and extrapolation performance on held-out temporal stages in retinal progenitor cell (RPC) data. **a**, Joint UMAP visualization comparing experimentally measured (grey) and predicted (colored) cell states across held-out test somite stages. Top rows depict interpolation (Somite stages 12, 15, 18), while bottom rows depict extrapolation (Somite stages 33, 34). For each panel, 1,000 measured and 1,000 predicted cells were sampled and projected into a shared embedding. CellPace predictions (red) and baseline predictions (blue) are overlaid on measured data to visually assess manifold alignment. **b**, Quantitative performance matrix evaluating 2-Wasserstein distance in PCA space (W2; lower is better) and miLISI (higher is better) for each test stage. Rows are ordered by average W2 performance. Circle background colors represent the rank of each method within the column (darker indicates better rank), while annotated numbers display the raw metric values. **c**, Statistical property distributions comparing measured and generated data in the five testing stages. Violin plots display cell-level metrics (zero fraction, library size, TMM normalization factor, expression variance, mean expression) and gene-level metrics (mean expression, zero fraction, library size). The scatter plot (bottom right) visualizes the relationship between per-cell zero fraction and library size. **d**, Distributional fidelity assessment via Energy distance. The heatmap compares the similarity between measured and generated distributions for the eight properties shown in **c**. Cell colors indicate the rank per metric (1/yellow = best, 7/dark = worst), while annotated values show the raw Energy distance. Methods are ordered by the mean Energy distance across all metrics, quantified in the right-hand bar plot (mean ± std).

We also evaluated each model based on its general data properties in the five testing stages (Fig. 3c-3d). In general, most methods broadly matched the real per-cell distributions of sparsity, library size and normalization factors, but several deviations were observed. scDiffusion and ESCFD displayed long-tailed distributions for Trimmed Mean of M-values (TMM) normalization factors and per-cell expression variance, with occasional cells showing extreme values that were absent in the real data, suggesting increased variability in library composition and high-variance gene representation. Similar patterns appeared at the gene level: for several baselines, per-gene mean expression and variance exhibited heavy tails, whereas the real data and CellPace-generated data remained more tightly concentrated. These discrepancies also appeared in the relationship between library size and zero fraction. Real cells showed a tight coupling between sequencing depth and sparsity, which CellPace closely reproduced, while scDiffusion and ESCFD generated more dispersed patterns that partially decoupled depth from sparsity. The energy-distance summary in Fig. 3d captures these trends quantitatively.

### CellPace preserves marker activation patterns and spatial positional information

Next, we set out to determine whether CellPace preserves higher-order organization in a complex, multi-lineage setting. We focused on the posterior embryo dataset, analyzed in detail in a previous study (Fig. 2 of [15]), comprised of neuromesodermal progenitors (NMPs), mesodermal progenitors (*Tbx6*⁺), notochord, gut, and ciliated nodal cells. Following the approach used in previous sections, we generated CellPace data and annotated cell types via k-nearest-neighbor label transfer (Methods, Supplementary Fig. 4a-4b).

**Fig. 4.**
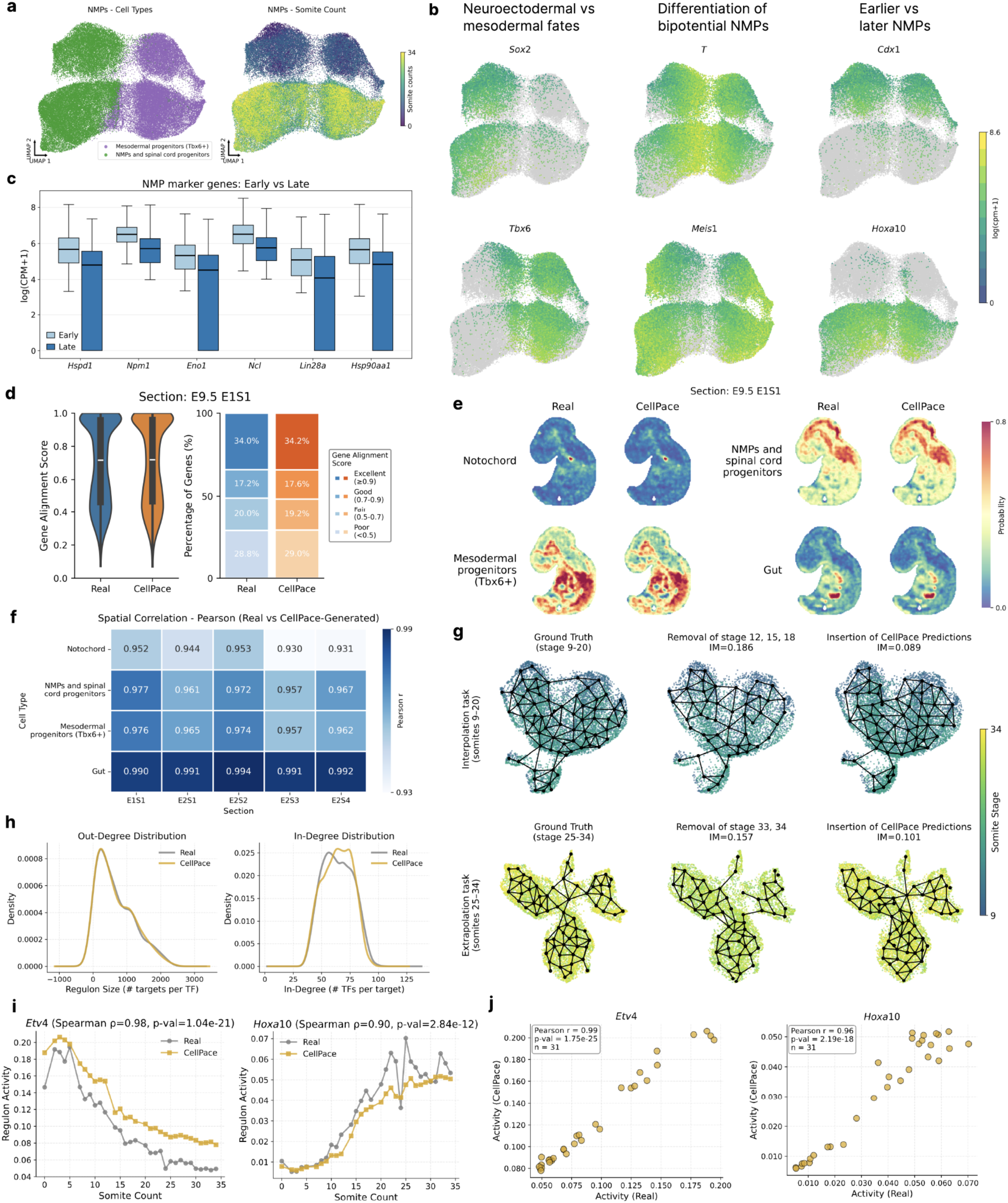
Analysis of biological signals in CellPace-generated posterior embryo samples. **a**, UMAP visualization of CellPace-generated neuromesodermal progenitors (NMPs) and mesodermal progenitors. Left: Cells colored by inferred cell identity, showing the separation of NMP/spinal cord progenitors (green) and mesodermal progenitors (purple). Right: Cells colored by somite counts, illustrating the continuous developmental dynamics from early (purple) to late (yellow) stages. **b**, Marker expression plots showing log-normalized expression of key regulatory genes, organized by biological function (columns). Left column: Complementary expression patterns of neuroectodermal (*Sox2*) versus mesodermal (*Tbx6*) fates. Middle column: Distinction between the bipotential NMP state (*T*) and differentiated progenitors (*Meis1*). Right column: Spatially distinct expression patterns corresponding to early (*Cdx1*) versus late (*Hoxa10*) temporal identity. **c**, Box plots of expression levels for metabolic and maintenance genes. Analysis is restricted to generated cells matching the specific NMP molecular signature, defined in [15]. Comparison groups are defined by developmental stage: Early (before somite 14) versus Late (somite 14 and after). Boxes denote the interquartile range (IQR), center lines mark medians, and whiskers extend to 1.5xIQR. **d**, Evaluation of single cell mapping to the Mouse Organogenesis Spatiotemporal Transcriptomic Atlas (MOSTA) using Tangram. Violin plots (left) and stacked bar charts (right) summarize the distribution of gene alignment scores for real and CellPace-generated scRNA-seq profiles mapped to spatial transcriptomics spots in the E9.5 embryo section E1S1. **e**, Spatial localization of predicted cell populations. Spatial probability maps for representative lineages (notochord, neuromesodermal progenitors and spinal cord progenitors, mesodermal progenitors, and gut) shown for section E1S1. Maps are obtained by mapping real (left) and CellPace-generated (right) scRNA-seq profiles to MOSTA spatial transcriptomics data using Tangram. **f**, Quantitative comparison of spatial probability maps. Heatmap showing Pearson correlation coefficients between spatial probability maps derived from real and CellPace-generated data, computed for four cell types across five anatomical sections (E1S1, E2S1-E2S4). **g**, Trajectory recovery analysis using PAGA graphs. Top: Interpolation task (Somite stages 9-20). Bottom: Extrapolation task. Graphs compare the topology of Ground Truth (left), the incomplete reference (middle), and the trajectory after inserting CellPace predictions (right). Structural dissimilarity is quantified using the Ipsen-Mikhailov (IM) distance, where lower values indicate greater topological similarity to the ground truth. **h**, Network topology comparison of gene regulatory networks (GRNs) inferred from real (grey) and CellPace-generated (gold) data. Curves show the kernel density estimates for out-degree (targets per TF) and in-degree (TFs per target) distributions. **i**, Temporal dynamics of key regulons. Line plots show the mean regulon activity of *Etv4* and *Hoxa10* across somite stages for real (grey) and CellPace-generated (gold) data. **j**, Comparison of regulon activity levels. Scatter plots showing the mean activity of *Etv4* and *Hoxa10* for real (x-axis) and CellPace (y-axis) data across all somite stages.

First, we resolved the fine-grained structure of the NMP lineage and its derivatives (Fig. 4a), which showed a similar developmental pattern to that observed in the original experimental data (compare with Fig. 2c in [15]). Projecting the expression of canonical marker genes onto this manifold showed that CellPace captures the divergence of neural (Sox2⁺) and mesodermal (Tbx6⁺) fates while preserving a transcriptional signature of bipotency, marked by high *T* (Brachyury) and low *Meis1* expression (Fig. 4b, compare with Fig. 2d in [15]). The model also captured the temporal “trunk-to-tail” transition within this lineage, reproducing the shift from *Cdx1* expression in early NMPs to *Hoxa10* in late NMPs (Fig. 4b). We quantified this temporal fidelity by comparing expression distributions of early-stage markers: CellPace-generated late NMPs showed the expected downregulation of genes such as *Hspd1*, *Npm1* and *Lin28a* compared with their early counterparts (Fig. 4c), consistent with the early-enriched module identified in the reference atlas (Fig. 2l right in [15]). We extended this analysis to the gut endoderm lineage, where re-embedding revealed a structured developmental continuum (Supplementary Fig. 4c). Marker gene analysis showed that CellPace resolves spatially distinct subpopulations along the anterior-posterior axis, localizing foregut/lung (*Nkx2-1*), liver (*Hhex*, *Afp*) and pancreas (*Onecut2*) identities to the anterior domain, while restricting *Cdx2* and *Hoxc8* expression to the posterior hindgut. This reconstruction of regionally restricted expression programmes suggests that CellPace captures positional information encoded within the single-cell transcriptome.

To further assess whether CellPace-generated cells preserve spatial positional information, we mapped both real and generated single-cell profiles to E9.5 mouse embryo sections from the Mouse Organogenesis Spatiotemporal Transcriptomic Atlas (MOSTA [20]) using Tangram [21]. Across five anatomical sections (E1S1, E2S1-E2S4), the distributions of per-gene alignment scores were highly concordant between real and CellPace-generated data (Fig. 4d; Supplementary Fig. 5a), indicating comparable mapping quality across sections. Spatial probability maps show that generated cells localize to anatomically distinct regions corresponding to their annotated lineages, closely matching the spatial patterns observed in real data (Fig. 4e; Supplementary Fig. 5b). At the spatial spot level, Tangram-derived cell-type enrichment scores exhibited strong quantitative agreement between real and generated profiles (Fig. 4f; Supplementary Fig. 6a). For all examined sections and lineages, enrichment values were highly correlated (Pearson r ≥ 0.93; Fig. 4f), with consistent rank-order preservation confirmed by Spearman correlations (Supplementary Fig. 6b).

### CellPace captures data manifold, developmental trajectory and regulatory signals

We next evaluated the ability of CellPace in reconstructing the data manifold and its topology in unseen stages. We first used PAGA [21] to reconstruct a graph pertaining to the underlying data topology and developmental trajectories using the experimental data in a window with ten measured stages, which included the unseen interpolation timepoints (Fig. 4g, top-left). We repeated the same analysis using stages that excluded the interpolation timepoints (Fig. 4g, top-middle) and using stages that replaced the interpolation timepoints with those generated by CellPace (Fig. 4g, top-right). Visually, incorporating CellPace-generated data improved the reconstructed graph, making it more similar to the graph obtained from the full real dataset. This observation was confirmed quantitatively: the Ipsen-Mikhailov (IM) distance [22] between the CellPace-augmented graph and the full-data graph was 0.089, compared with 0.186 for the ablated dataset. We repeated this analysis for another window covering the extrapolation stages, which resulted in similar conclusions: both visually (Fig. 4g, bottom) and quantitatively, the CellPace-augmented graph improved similarity to the full-data graph (IM = 0.101) compared to the ablated-data graph (IM = 0.157).

Finally, we assessed whether CellPace captures network-level relationships of genes in the form of gene regulatory networks (GRNs) and their temporal activity during developmental transitions. We inferred gene regulatory networks (GRNs) from the real and CellPace-generated datasets separately using pySCENIC [23] and compared their global topological properties. Overall, CellPace reproduced key features of the network architecture, with out-degree (“regulon size”) and in-degree (“regulatory convergence”) distributions broadly matching those of the real data (Fig. 4h). To assess temporal fidelity, we first visualized regulon activity profiles across somite stages for representative regulators. CellPace recapitulated the expected opposing programs in the posterior growth zone (Fig. 4i), including the progressive downregulation of the FGF effector *Etv4* (Spearman ρ = 0.98) and the increasing activity of the NMP-associated factor *Hoxa10* (Spearman ρ = 0.90). We then quantified agreement by correlating regulon activity between real and generated data across the 68 shared regulons, showing a high degree of concordance (median Pearson r = 0.81, Supplementary Table 3). This trend is illustrated by the strong linear correspondence for exemplar drivers when comparing stage-wise mean regulon activity (Fig. 4j): *Etv4* (Pearson r = 0.99) and *Hoxa10* (Pearson r = 0.96).

### A multimodal extension of CellPace enables joint modeling of transcriptomics and chromatin accessibility dynamics in palate development

Given that the TDiF architecture operates in the latent space, CellPace can be easily extended to enable multimodal generation using appropriate encoder/decoders. To illustrate this, we replaced the original VAE in CellPace (scVI [11]) with MultiVI [24] (Fig. 5a), a small modification that enables CellPace to jointly model and generate scRNA and scATAC data. We used a single-cell multiome dataset of mouse secondary palate development (E12.5 to E14.5), containing measurements of n = 36,154 cells [25]. To facilitate fine-grained temporal analysis, we used pseudotime obtained using CellRank2 [4] followed by palantir [26] and binned cells into 50 intervals (Methods). We held out eight bins (5, 10, 15, 20, 25, 30, 35, 40) for interpolation analysis and the last four bins (46 to 49) for extrapolation analysis.

**Fig. 5.**
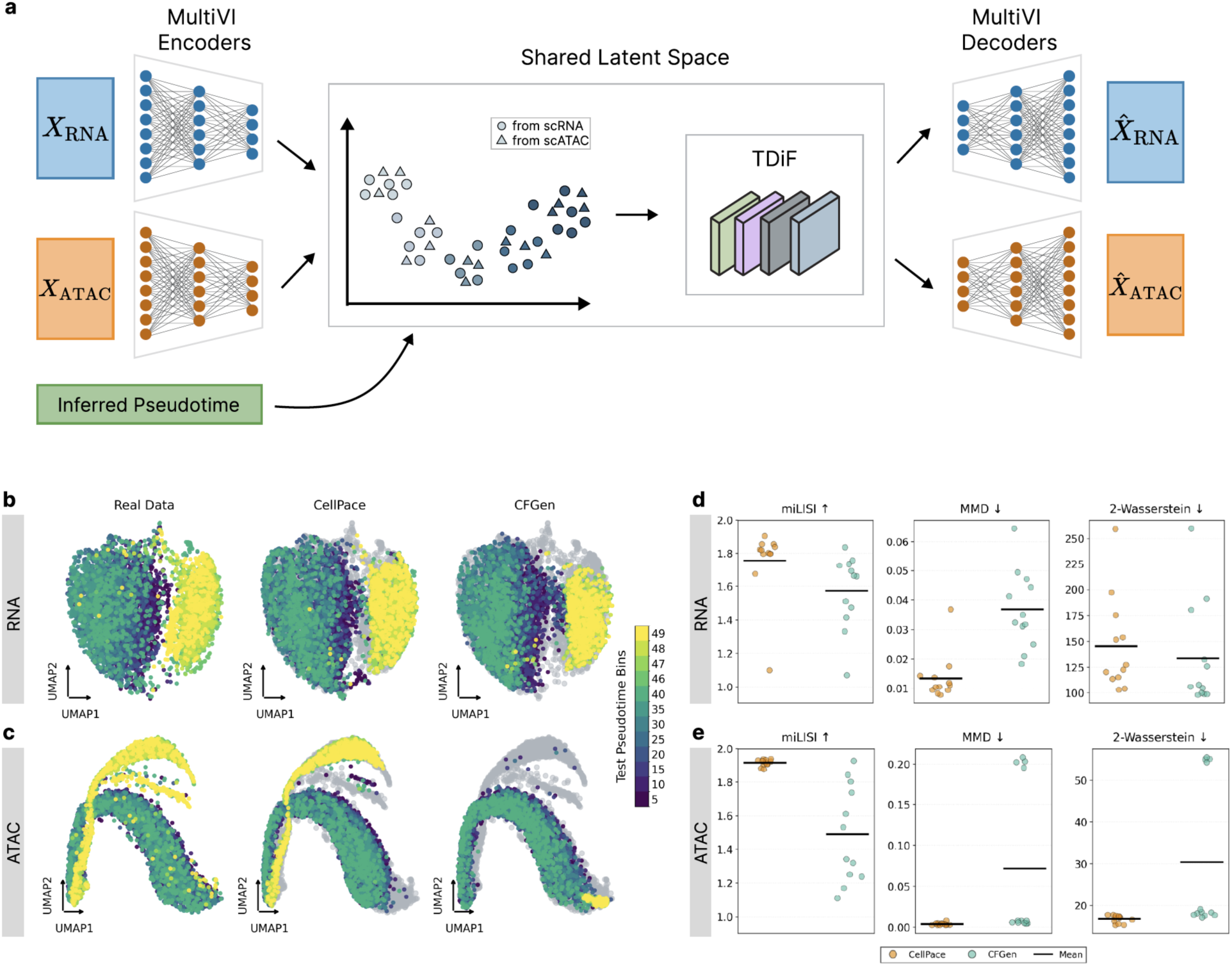
Multimodal temporal generation of the mouse palate data. **a**, Multimodal generation framework of CellPace. Paired scRNA-seq and scATAC-seq profiles are encoded into a shared latent space via MultiVI. The TDiF module denoises latent sequences conditioned on pseudotime, and modality-specific decoders generate aligned gene expression (ZINB distribution) and chromatin accessibility (Bernoulli distribution) profiles from the denoised latent representations. **b**-**c**, Temporal manifold visualization. UMAP embeddings of scRNA-seq (**b**) and scATAC-seq (**c**) profiles. Columns compare real data (left) with generated samples from CellPace (center) and CFGen (right). Generated cells (colored by pseudotime bins) are overlaid onto the real manifold (grey background) to visualize temporal alignment in the shared embedding space. **d**-**e**, Quantitative assessment of data generation. Box plots summarizing generation quality across held-out temporal stages (n = 12), including both interpolation and extrapolation settings for RNA (**d**) and ATAC (**e**). Metrics include miLISI (higher is better), Maximum Mean Discrepancy (MMD; lower is better), and 2-Wasserstein distance (lower is better).

Since CFGen is the only model in our cohort that supports multimodal generation, we used it as our baseline. Visually, both CellPace (multimodal) and CFGen (multimodal) performed well on the transcriptomic data (Fig. 5b), with extrapolation being the more challenging task. For scATAC generation, both CellPace and CFGen performed relatively well in the interpolation stages, but CFGen struggled in the extrapolation task (Fig. 5c). The real scATAC data displayed a multi-branched architecture representing diverging late-stage lineages. CellPace largely recovered this structure, populating the main path and the lower branch, although the intermediate secondary branch is only weakly represented (Fig. 5c). CFGen failed to populate the upper branches altogether, yielding a truncated manifold in which distinct late-stage lineages collapse into a single diffuse cloud.

These observations were confirmed quantitatively. Both models achieved strong performance in transcriptomic generation (Fig. 5d). In generating scATAC data, however, CellPace outperformed CFGen in all metrics (Fig. 5e). Particularly in the extrapolation task, CellPace obtained mean miLISI and MMD of 1.85 and 3.6E-3, respectively, compared to CFGen with mean miLISI = 1.37 and mean MMD=5.2E-3. These analyses demonstrate the flexibility of the CellPace framework for generating multimodal data. Moreover, CellPace’s ability to generate realistic data using pseudotime underscores its broad utility, even for snapshot datasets lacking explicit temporal measurements.

## Discussion

CellPace is designed to bridge the gap between cross-sectional single-cell datasets and the continuous developmental processes they sample. Unlike existing generative models that treat developmental stages as discrete, independent categories, CellPace learns a continuous-time prior, enabling the generation of *de novo* cell populations at arbitrary developmental positions. Across four mouse lineages, CellPace reproduced the distribution and temporal organization of observed stages and maintained high fidelity when interpolating or extrapolating to held-out somite stages. In the more complex posterior embryo, it preserved multi-lineage structure, gene-regulatory programs, and spatial patterns. In a mouse palate 10x multiome dataset, CellPace extended naturally to paired RNA-ATAC generation, even when the temporal axis had to be reconstructed from pseudotime rather than explicit staging.

Our analyses indicate that CellPace retains biologically meaningful structure at several levels. In the posterior embryo, the model recovered known developmental features, including the separation of neuromesodermal progenitors into neural and mesodermal branches and the anterior-posterior patterning of gut progenitors. Spatial mapping to the MOSTA atlas further showed that CellPace-generated cells retain sufficient positional information to be assigned to anatomically specific regions with confidence comparable to real cells. Gene-regulatory networks inferred from generated data captured the degree distributions and stage-dependent activity profiles of key regulons. This indicates that the diffusion dynamics preserve not only marginal gene expression but also coordinated regulatory programs that drive lineage commitment. In the multiome setting, the model captured joint RNA-ATAC correlations at held-out late pseudotime bins, a regime where the alternative model struggled. This capacity to model joint dynamics from inferred pseudotime underscores the framework’s robustness to sparse or irregular temporal sampling, a common challenge in atlas-scale integration where dense experimental time series are often unavailable.

In our analyses, we observed some models (particularly CFGen) to perform well during simulation based on observed timepoints, while struggling in interpolation or temporal forecasting. One should note that CFGen and other similar models are designed for class-conditional generation and do not naturally support temporal extrapolation. Our post-hoc approach represents a best-effort, user-level adaptation, rather than a task-specific redesign. As such, their performance in these regimes should be considered a practical reference point rather than a definitive statement about each method’s theoretical limits. However, this further underscores the necessity of frameworks like CellPace that explicitly model temporal continuity for predictive tasks.

We believe CellPace represents a substantive step toward a more general family of temporal generative models for single-cell biology. One natural extension is to move beyond transcriptomic and chromatin accessibility to multi-layer representations in which additionally, genomic information, regulatory elements, and protein features are mapped into the shared latent space before temporal diffusion, enabling a unified treatment of the constraints on cell state. A second direction is to incorporate spatially resolved measurements so that CellPace models continuous spatiotemporal fields, learning not only how cell states change over time but also how they organize within developing tissues. Finally, the same architecture invites cross-species transfer learning: by aligning latent manifolds across species, one could train dynamics where core developmental programs are shared while species-specific adapters capture divergent trajectories. Together, these extensions point toward a more general, sequence-aware, and spatially grounded representation of cellular dynamics built on the foundations established here.

## Methods

### The architecture of CellPace

CellPace is a generative framework designed to model the temporal dynamics of single-cell omics data. First, a VAE is trained to learn low-dimensional latent representations of individual cells, effectively capturing the manifold of cellular identity. Given this pretrained VAE, a Transformer-based diffusion model called TDiF (Temporal Diffusion Forcing) learns the transition probabilities between these latent states over time (Fig. 1a-1d).

#### Latent Representation Learning using VAE

To avoid the sparsity and high dimensionality of raw count data, we first encode cells into a compressed latent space. For single-modality scRNA-seq datasets, we used scVI [11] to learn latent representations, modeling sparse and over-dispersed gene counts with a Zero-Inflated Negative Binomial (ZINB) likelihood. For paired multiome datasets, we utilized MultiVI [24] to learn a joint latent representation that integrates information from both RNA and chromatin modalities.

#### Temporal Diffusion Forcing (TDiF)

To model cellular transitions within this latent space, we designed an architecture capable of capturing continuous temporal dynamics without the structural limitations of existing generative backbones. Standard U-Nets [27, 28], common in image diffusion, rely on convolutional down sampling and spatial locality—assumptions ill-suited for unordered gene sets. Moreover, their bidirectional processing allows later states to influence the reconstruction of earlier ones during training, violating temporal causality. While recurrent neural networks enforce directionality, they often suffer from vanishing gradients [29] over long developmental trajectories. Similarly, while token-based Transformers [30] capture long-range dependencies, they typically require quantizing continuous molecular profiles into categorical vocabularies, which can obscure the fine-grained metric geometry of the cellular manifold.

To overcome these challenges, we developed Temporal Diffusion Forcing (TDiF), a backbone motivated by the Diffusion Forcing (DiF) [10] framework. Unlike traditional diffusion models that denoise entire sequences at a single noise level, TDiF uses per-position noise levels as a continuous masking signal. This design allows low-noise past frames to act as known context while high-noise future frames serve as uncertain targets within a single forward pass, enabling flexible, causal sequence generation (Fig. 1b-1d). More importantly, its gap-aware temporal encoding enables capturing of continuous temporal dynamics.

#### Gap-Aware Temporal Encoding

A critical innovation in TDiF is its handling of time. Sequence diffusion models typically encode temporal structure using sinusoidal embeddings of integer indices, e.g., x = x + PositionalEmbed (0, 1, …, T − 1), implicitly assuming uniformly spaced observations. In developmental biology, sampling is often irregular and the time gaps between stages (for example, between stages with somite counts of 10 and 18) are biologically informative. Thus, in TDiF we replaced these positional embeddings with a continuous biological time embedding that explicitly encodes both position along the developmental axis and the temporal spacing between observations. Specifically, for each developmental state s_t_, we constructed a two-dimensional temporal feature vector h_t_ = [τ_t_, Δ_t_] consisting of:

(1) normalized developmental position 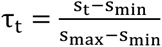, which maps the developmental stage s_t_ to the interval [0, 1] using the minimum and maximum stages observed during training and enables the model to generalize to previously unseen stages within or beyond this range, and
(2) elapsed time gap Δ_t_ = s_t_ − s_t–1_, which captures the temporal interval since the previous position in the sequence (with Δ_1_ = 0 for the first position).

By providing explicit gap information, CellPace can learn distinct denoising dynamics for short-range transitions (for example, consecutive somite stages with Δ = 1) versus long-range developmental leaps (for example, interpolating across a gap of Δ ≥ 5). The gap features are linearly scaled before processing to maintain numerical stability.

To inject this biological context, we used a Transformer backbone equipped with AdaLN (Fig. 1d). Rather than concatenating time features only at the input, the vector h_t_ is passed through a small multilayer perceptron (MLP) to produce scale and shift parameters [scale_t_, shift_t_] = MLP(h_t_), which modulate activations within every Transformer block:

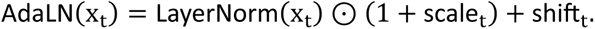

The input pipeline concatenates noised latent embeddings 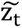 with sinusoidal embeddings of the noise levels k_t_, and the resulting features are processed by the AdaLN-modulated Transformer. We used AdaLN-Zero initialization for the temporal conditioning MLP (final layer initialized to zero), so that blocks behave as standard LayerNorm at the start of training and temporal modulation is introduced gradually.

### Training and Optimization of CellPace

CellPace training follows a two-stage procedure. In Stage 1, we train the VAE (scVI or MultiVI) to maximize the ELBO. In Stage 2, we train the TDiF model on sequences of VAE latent representations. We construct a training dataset by sampling temporal sequences using a geometric strategy that balances local and long-range dependencies. For each sequence, we randomly select a starting stage, then iteratively sample subsequent stages by drawing jumps from a geometric distribution whose parameter is chosen so that the expected gap matches the mean stage gap observed in the training data. For each stage in the sampled sequence, we randomly select a cell from that stage’s latent representations and compute its gap-aware temporal features h_t_. This sampling strategy ensures the model encounters both consecutive stage transitions and larger developmental leaps during training, enabling it to learn denoising dynamics across multiple temporal scales. The TDiF model is trained on sequences sampled from a static dataset of latent vector representations, which are generated by a one-time deterministic encoding of the real cells using the frozen VAE’s posterior mean.

We sample diffusion noise levels k_t_ independently for each temporal position (from a uniform distribution of timesteps) rather than sharing a global noise level across the sequence. This uncorrelated sampling strategy forces the network to learn the denoising vector field across all combinations of uncertainty simultaneously, effectively enabling the “continuous masking” mechanism required for flexible causal generation. The model is then trained to predict the original clean latent by minimizing the following weighted mean-squared error objective:

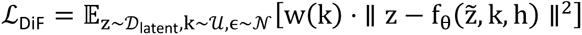

where 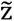 is the noised latent sequence, obtained via the standard forward process of the Denoising Diffusion Probabilistic Model 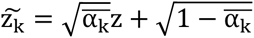, and h contains the gap-aware temporal features. The weight function w(k) applies the min-SNR weighting strategy [13]. This weight, derived from the signal-to-noise ratio 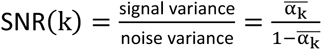, is clipped at a constant threshold. This prevents the overall loss from being dominated by high-SNR (near-clean) samples, which have low error but contribute disproportionately to the gradient, and balances the optimization across the full spectrum of noise levels.

For parameters and implementation details see Supplementary Note 1.

### Generation of new samples from noise

To generate new sequences of cell states across developmental time, we use an autoregressive procedure called pyramid scheduling [10] that synthesizes entire sequences of latent cell states (Fig. 1b-1c). For a given starting point and target horizon, latent representations at all future timepoints are initialized as Gaussian noise and denoised in a cascading pattern: earlier timepoints receive more denoising steps compared to later ones, allowing uncertainty to propagate forward in time. To extrapolate beyond the training horizon, we use a sliding-window scheme: the first step synthesizes a full block, matching the training sequence length (e.g., five stages), and subsequent steps advance the window by a chosen chunk size using previously generated latents as context.

Inference is accelerated using DDIM sampling; standard DDPM sampling is also supported when the number of sampling steps matches the training schedule. This combination of pyramid scheduling, which allocates denoising effort across timepoints, and DDIM, which accelerates latent updates, enables efficient generation of coherent developmental sequences of latent cell states. Once clean latent representations z are obtained, we decode them to gene expression using the frozen VAE decoder. For multi-omics models trained with MultiVI, we generate the gene-expression modality from the joint latent representation and sample chromatin accessibility separately from its Bernoulli output distribution.

### Datasets and Preprocessing

#### Single-cell RNA-seq datasets

We obtained developmental single-cell RNA-seq data from the mouse embryo atlas [15], which profiles over 11 million cells across embryonic days E6.5-E13.5. To focus on lineages with fine-grained temporal resolution, we restricted our analysis to cells with somite-level stage annotations, excluding cells labelled “somite_na” (approximately 2 million cells across 32 somite stages), and to protein-coding genes. From this subset, we selected three progenitor cell types for temporal modelling based on two criteria: (1) sufficient temporal coverage, with at least 50 cells per somite stage for most stages to enable stable train-test splits; and (2) representation of diverse developmental contexts and abundance levels. The resulting datasets comprised epithelial cells (n = 54k), retinal progenitor cells (n = 35k) and embryonic blood vessel endothelial progenitor cells (n = 9k).

To assess whether CellPace can preserve biological heterogeneity when modelling multiple interacting lineages simultaneously, we additionally constructed a fourth dataset, “posterior embryo” (n = 121k), comprising five spatially co-located but transcriptionally distinct groups: notochord, ciliated nodal cells, neuromesodermal progenitors (NMPs) and spinal cord progenitors (combined), gut, and mesodermal progenitors (embryonic days E8.0-E10.0), following the filtering criteria in [15].

All datasets underwent standardized preprocessing using Scanpy v1.11 [31]: quality-control filtering (minimum 100 genes per cell and minimum 3 cells per gene), doublet detection using Scrublet [32], normalization to 10,000 counts per cell and log₁p transformation. Raw count matrices were retained in the AnnData object (layers[“raw_counts”]) and used directly for model training.

For each scRNA-seq dataset, we employed a stage-based holdout strategy to evaluate temporal generalization. Because the datasets provide only discrete somite stages, simple random splitting would ignore differences in biological complexity across time. To construct a more informative test set, we first quantified the “difficulty” of generating each somite in PCA space using two metrics: intra-somite heterogeneity (coefficient of variation of pairwise distances) and temporal dynamics (maximum mean discrepancy, MMD, to the preceding somite) (Supplementary Fig. 7). Based on this stratification, we defined a deterministic split, designed to stress-test specific modelling capabilities while respecting the temporal structure of the series.

To form the test set, interpolation targets were chosen to span the difficulty spectrum: somite 18 (“easiest”; low heterogeneity/low velocity), somite 15 (“moderate”) and somite 12 (“hardest”; high heterogeneity/high velocity). For extrapolation, the terminal timepoints, somite stages 33 and 34, were used as future-state test targets. To form the validation set, which was used exclusively for hyperparameter optimization, we selected S10, S11, S22, S31 and S32. The remaining somite stages were selected as the training set.

To avoid data leakage, we derived feature-selection and normalization statistics exclusively from training stages. We applied Scanpy’s Seurat-flavoured HVG selection [31] on training somite stages only (3,000 genes) and then used this fixed gene set for validation and test cells. For the posterior embryo dataset, we additionally included lineage-specific marker genes highlighted in [15] (for example, *Sox2, Cdx2, Sox17, Msgn1, Foxj1 and Wnt* pathway genes) to retain key developmental regulators even when they were not among the top HVGs. This procedure ensured that no information from held-out stages influenced gene selection, normalization, parameters or other preprocessing decisions, maintaining a clear separation between training and evaluation data throughout the pipeline.

#### Paired RNA-ATAC 10x multiome dataset

We obtained a mouse palate development multiome dataset (n = 36k) from [25] via the public repository (https://github.com/fangfang0906/Single_cell_multiome_palate). Each modality was processed separately before integration. For RNA, we applied Scanpy’s Seurat-flavoured HVG selection to retain 3,000 HVG. For ATAC, we filtered peaks to those present in at least 5% of cells to obtain a feature set compatible with variational autoencoder training. Preprocessing and trajectory inference were performed on log-normalized data, and the raw integer counts of these filtered features were retained in the final MuData object for model training.

We used trajectory inference to define a temporal axis. We applied CellRank2 [4] with the RealTimeKernel, computing optimal transport with MOSCOT and constructing a cell-state transition matrix that combined RNA neighbour connectivity (20% weight) and optimal transport distances (80% weight). We then symmetrized this transition matrix to satisfy the undirected-graph requirement of diffusion maps. The root cell was identified automatically as the cell with the maximum value in the first diffusion component (DC1) within the E12.5 timepoint. We computed pseudotime using Palantir on the symmetrized transition matrix and binned it into 50 equal-frequency intervals by quantile binning, yielding approximately balanced cell counts per stage (labels: pt_00 to pt_49).

To evaluate temporal generalization, we applied a deterministic uniform-interval splitting strategy to partition the 50 bins into training and test sets. Interpolation targets were defined by holding out every fifth bin across the trajectory (pt_05, pt_10, pt_15, pt_20, pt_25, pt_30, pt_35, pt_40). For extrapolation, the terminal segment of the trajectory was held out as future-state test targets (pt_46-pt_49).

### Baseline models for temporal scRNA-seq generation

To assess CellPace against existing approaches for temporal single-cell generation, we benchmarked it against a panel of recently proposed generative models spanning diffusion, flow matching and ODE/SDE-based methods. scDiffusion [8] is a conditional latent diffusion model for generating single-cell RNA-seq profiles with specified cellular attributes that utilizes classifier guidance. cfDiffusion [9] is a two-stage conditional latent diffusion model that reuses the deterministic autoencoder backbone of scDiffusion but replaces the separate noise-conditioned classifier with classifier-free guidance. ESCFD [18] is a three-stage latent diffusion framework that combines an autoencoder, a masked autoencoder and a diffusion Transformer to model conditional single-cell RNA-seq distributions. CFGen [5] is a two-stage conditional generative model that combines a deterministic autoencoder with flow matching in a learned latent space to model stage-conditional single-cell RNA-seq distributions. scNODE [6] is a conditional generative model that combines a variational autoencoder with neural ordinary differential equations to model developmental trajectories in continuous time. scIMF [7] is an SDE-based model that extends trajectory-based single-cell models by incorporating explicit cell-cell interactions through a McKean-Vlasov stochastic differential equation. A detailed description of these models is provided in the Supplementary Note 2.

All baselines were trained on the same developmental scRNA-seq datasets, using identical somite-based train/validation/test splits and the same set of HVG as CellPace for each processed dataset. The benchmarking pipeline followed a common pattern: for each method and dataset, we (i) converted the preprocessed AnnData objects to the method-specific input format, (ii) trained the generative model on training somite stages and (iii) generated 1,000 cells for each target somite stage. We saved generated cells as AnnData objects for evaluation.

### Multimodal generation using CFGen

Within our benchmark suite, CFGen [5] is the only method with native multimodal support, enabling joint RNA-ATAC generation. CFGen offers two multimodal configurations. In the default configuration, RNA and ATAC are each encoded into separate 100-dimensional latent spaces, and these embeddings are concatenated into a 200-dimensional vector that is passed directly to the flow-matching backbone. In an optional joint-latent configuration, the concatenated 200-dimensional vector is further mapped through a small multilayer perceptron to produce a shared 100-dimensional latent representation.

In our experiments, we used this joint-latent configuration. This choice not only aligns CFGen with the MultiVI backbone used in CellPace, where RNA and ATAC are embedded into a shared latent space, but also reflects the structure of our 10x multiome data, which provide paired RNA-ATAC profiles rather than unpaired modalities. After encoding, the flow-matching backbone operated on the 100-dimensional joint latent space, and the decoder remained split by modality, using a negative binomial likelihood for RNA counts and a Bernoulli likelihood for ATAC peak accessibility. The total loss was the sum of RNA and ATAC reconstruction terms. Conditional generation used the same stage-conditional flow-matching backbone: joint latent samples were decoded into paired RNA and ATAC profiles, allowing CFGen to generate multimodal single cells conditioned on developmental stage or other categorical covariates.

### Evaluation Metrics

miLISI (median Local Inverse Simpson Index) quantifies mixing of “real” and “generated” data by first computing a joint t-SNE embedding (perplexity=30), and then measuring local diversity within k-nearest neighbor graphs (k=90, derived from 3×perplexity) using an adaptive Gaussian kernel with perplexity-based bandwidth optimization; higher miLISI values (approaching 2.0) indicate better mixing between real and generated cells and a higher quality of data generation.

2-Wasserstein distance measures distributional divergence using optimal transport, where we first compute a joint PCA on the concatenated real and generated data (selecting min(50, n_samples-1) components), then split the transformed coordinates back to separate populations and compute exact earth mover’s distance with squared Euclidean cost matrix (power=2); lower values indicate better distributional matching.

1-Wasserstein distance follows the same joint PCA projection strategy but uses linear cost (power=1), providing a metric more robust to outliers.

MMD (Maximum Mean Discrepancy) employs a multi-scale Gaussian kernel with data-adaptive bandwidths determined by the median heuristic: we estimate the median distance among the 25 nearest neighbors across 20 bootstrap samples of the real data, then construct three kernel scales at 0.5×median, 1×median, and 2×median with equal weights (one third each), yielding a distribution-free test statistic that captures differences across multiple resolutions.

For all metrics, when the number of generated cells differs from the number of real cells at a given stage, we randomly subsampled both groups to the smaller size using a fixed random seed. For the palate RNA-ATAC multiome experiments, we applied the metrics to the real and generated gene expression data using the interpolation and extrapolation bin splits defined in Paired RNA-ATAC 10x multiome dataset.

### Trajectory-based evaluation

To assess how well generated cells recapitulate developmental dynamics, we employed a reference-based trajectory inference approach where pseudotime is first computed on real data using either Diffusion Pseudotime (DPT) [19] or Palantir [26]. Generated cells were then projected into the real data’s PCA space (50 PC components fitted on real training cells), and pseudotime values were transferred via k-nearest neighbor regression (k = 30) to ensure all methods share a common pseudotime scale.

### Cell-type and stage-resolved Expression Patterns in posterior embryo

To validate the preservation of biological structure, we harmonized real and CellPace-generated datasets (somite stages 0-34) to a shared gene set and capped each somite at 4,000 cells. We then normalized and log-transformed the data before dimensionality reduction. We performed a joint principal component analysis (PCA; 30 components) on the concatenated dataset and transferred cell-type annotations from real to generated cells via k-nearest neighbour classification (k = 5) in this joint PCA space. We visualized global topology using a 3D UMAP embedding (30 neighbours, min_dist = 0.3).

For lineage-specific analysis, we re-embedded NMP/mesoderm and gut subsets in two dimensions to resolve substructure. We visualized marker gene expression for cell fate specification (*Sox2*, *Tbx6*), NMP identity (*T, Meis1*) and gut regionalization (*Nkx2-1, Cdx2*) using log-normalized counts. To quantify developmental timing signatures, we defined NMPs based on raw count thresholds (*T* ≥ 5 and *Meis1* = 0) following the setup in [15] and summarized expression distributions with box plots showing the median, interquartile (IQR) range and 1.5× IQR whiskers.

### Gene Regulatory Network Inference

We inferred gene regulatory networks (GRNs) independently from real and CellPace-generated posterior embryo data (pooled across somite stages 0-34) using the pySCENIC pipeline (v0.12.1) with mm10 motif databases (10 kb upstream and downstream of transcription start sites). The workflow comprised three steps: (1) inference of co-expression modules from raw count matrices using GRNBoost2 [33] (gradient boosting regression; 8 workers); (2) pruning of indirect targets via cisTarget [34] motif enrichment analysis to retain direct TF-target interactions; and (3) quantification of regulon activity per cell using AUCell. We characterized network topology by comparing in-degree (regulatory convergence) and out-degree (regulon size) distributions via kernel density estimation. We assessed developmental dynamics by aggregating mean AUCell scores per somite stage and quantified concordance between real and generated temporal profiles using Spearman’s rank correlation.

### PAGA-based Graph Topology Analysis

We used PAGA-based graph analysis to quantify how posterior embryo graph structure changes when developmental stages are removed and subsequently augmented with CellPace predictions, adapting the trajectory-topology comparison strategy introduced in a previous study [7]. The posterior embryo dataset exhibits irregular somite coverage, with somite stages 13 and 19 absent from the real data. To obtain balanced windows, we selected two ten-somite intervals with complete real coverage: an interpolation interval spanning somite stages 9-20 (10 observed stages: 9, 10, 11, 12, 14, 15, 16, 17, 18, 20; held-out test stages 12, 15 and 18 lie within this temporal window) and an extrapolation interval spanning somite stages 25-34 (10 observed stages: 25-34; held-out test stages 33 and 34 extend beyond the highest training stage). This design ensures that both windows contain exactly ten observed somite stages despite gaps in developmental stage annotation, providing comparable statistical power and visual balance for evaluating interpolation and extrapolation settings.

For each window, we subsampled 1,000 cells per somite, performed PCA (50 components) and fitted a UMAP embedding (50 neighbours, min_dist = 0.25) on the ground-truth dataset containing all real stages. We then used the same UMAP model to embed the ablated (training stages only) and CellPace-augmented (training stages plus CellPace predictions for held-out stages) datasets via the transform operation, ensuring that all conditions were represented in a shared low-dimensional space.

We applied Leiden clustering (resolution = 0.5) independently to each embedded dataset, allowing cluster counts to vary with data density and local structure. We constructed PAGA graphs from the resulting neighbour graphs using an edge-weight threshold of ≥ 0.1 and a minimum of 5 neighbours, yielding three graphs per window: (i) ground truth (all real stages), (ii) ablated (real training stages with held-out somite stages removed) and (iii) augmented (real training stages plus CellPace predictions for the held-out somite stages). We quantified topological similarity using the Ipsen-Mikhailov (IM) distance, a spectral metric that compares eigenvalue distributions of graph Laplacians without requiring explicit node matching (netrd v0.3.0).

### Spatial Mapping and Concordance Analysis

We mapped single-cell transcriptomes from real and CellPace-generated posterior embryos to five coronal tissue sections (E1S1, E2S1-E2S4) from embryonic day 9.5 of the Mouse Organogenesis Spatiotemporal Transcriptomic Atlas (MOSTA [20]) using Tangram [21]. We first harmonized datasets to the intersection of genes shared with the spatial atlas. Tangram was then trained to align single-cell expression profiles with spatial spot expression by minimizing a cosine-similarity loss, yielding per-spot cell-type probability distributions after projecting single-cell annotations onto spatial coordinates. We evaluated mapping quality using per-gene training scores, defined as the cosine similarity between predicted and measured spatial expression. We compared score distributions for real and CellPace-generated mappings using two-sided Kolmogorov-Smirnov tests and stratified genes into four quality tiers (Poor < 0.5; Fair 0.5-0.7; Good 0.7-0.9; Excellent > 0.9) for visualization.

To generate continuous spatial probability maps, we smoothed spot-level probabilities using k-nearest-neighbour averaging with k = round(log_2_(N_spots_)) + 1, followed by min-max normalization. We quantified spatial concordance between real and CellPace-generated distributions for notochord, neuromesodermal progenitors (NMPs) and spinal cord progenitors, *Tbx6*+ mesodermal progenitors and gut using Pearson and Spearman correlations between probability vectors across all spatial spots.

## Software and Data Availability

All datasets analyzed in this study are publicly available. The mouse embryo developmental scRNA-seq datasets were obtained from Qiu et al. via the CZ CellxGene Collection (Collection ID: 45d5d2c3-bc28-4814-aed6-0bb6f0e11c82). The processed mouse palate 10x multiome (RNA– ATAC) dataset was obtained from Yan et al. via their public repository https://github.com/fangfang0906/Single_cell_multiome_palate.

The CellPace software package is open-source and available on GitHub at https://github.com/Emad-COMBINE-lab/CellPace-release.

## Acknowledgements

This study was supported by a grant jointly funded by Natural Sciences and Engineering Research Council of Canada (NSERC) [ALLRP 586837 – 23] and FRQNT [2024-NOVA-344140] to AE and by a grant from NSERC [RGPIN-2019-04460] to AE. This work was also supported by resource allocations from Digital Research Alliance of Canada (alliancecan.ca) and Calcul Québec (www.calculquebec.ca) to AE. CS was supported by a PhD scholarship from FRQNT.

## Author Contributions

A.E. and C.S. conceived the project and contributed to manuscript writing. A.E. supervised the project and obtained funding. C.S. developed the framework and conducted all experiments and analyses. All authors read, revised and approved the final manuscript.

## Reference

1. Weinreb, C., et al., Fundamental limits on dynamic inference from single-cell snapshots. Proceedings of the National Academy of Sciences, 2018. 115(10): p. E2467–E2476.

2. Lähnemann, D., et al., Eleven grand challenges in single-cell data science. Genome biology, 2020. 21(1): p. 31.

3. Wolf, F.A., et al., PAGA: graph abstraction reconciles clustering with trajectory inference through a topology preserving map of single cells. Genome biology, 2019. 20(1): p. 59.

4. Weiler, P., et al., CellRank 2: unified fate mapping in multiview single-cell data. Nature Methods, 2024. 21(7): p. 1196–1205.

5. Palma, A., et al., Generating multi-modal and multi-attribute single-cell counts with CFGen. arXiv preprint arXiv:2407.11734, 2024.

6. Zhang, J., et al., scNODE: generative model for temporal single cell transcriptomic data prediction. Bioinformatics, 2024. 40(Supplement_2): p. ii146-ii154.

7. Jiang, Q., et al., Learning collective multi-cellular dynamics from temporal scRNA-seq via a transformer-enhanced Neural SDE. arXiv preprint arXiv:2505.16492, 2025.

8. Luo, E., et al., scDiffusion: conditional generation of high-quality single-cell data using diffusion model. Bioinformatics, 2024. 40(9): p. btae518.

9. Zhang, T., et al., cfDiffusion: diffusion-based efficient generation of high quality scRNA-seq data with classifier-free guidance. Briefings in Bioinformatics, 2025. 26(1): p. bbaf071.

10. Chen, B., et al., Diffusion forcing: Next-token prediction meets full-sequence diffusion. Advances in Neural Information Processing Systems, 2024. 37: p. 24081–24125.

11. Lopez, R., et al., Deep generative modeling for single-cell transcriptomics. Nature methods, 2018. 15(12): p. 1053–1058.

12. Peebles, W. and S. Xie. Scalable diffusion models with transformers. in Proceedings of the IEEE/CVF international conference on computer vision. 2023.

13. Hang, T., et al. Efficient diffusion training via min-snr weighting strategy. in Proceedings of the IEEE/CVF international conference on computer vision. 2023.

14. Song, J., C. Meng, and S. Ermon, Denoising diffusion implicit models. arXiv preprint arXiv:2010.02502, 2020.

15. Qiu, C., et al., A single-cell time-lapse of mouse prenatal development from gastrula to birth. Nature, 2024. 626(8001): p. 1084–1093.

16. Luo, H., et al., Forkhead box N4 (Foxn4) activates Dll4-Notch signaling to suppress photoreceptor cell fates of early retinal progenitors. Proceedings of the National Academy of Sciences, 2012. 109(9): p. E553–E562.

17. Xiao, D., K. Jin, and M. Xiang, Necessity and sufficiency of Ldb1 in the generation, differentiation and maintenance of non-photoreceptor cell types during retinal development. Frontiers in Molecular Neuroscience, 2018. 11: p. 271.

18. Li, S., et al. ESCFD: Probabilistic Flow Diffusion Model for Accelerated High-Quality Single-Cell RNA-seq Data Synthesis. in Proceedings of the 31st ACM SIGKDD Conference on Knowledge Discovery and Data Mining V. 2. 2025.

19. Haghverdi, L., et al., Diffusion pseudotime robustly reconstructs lineage branching. Nature methods, 2016. 13(10): p. 845–848.

20. Chen, A., et al., Spatiotemporal transcriptomic atlas of mouse organogenesis using DNA nanoball-patterned arrays. Cell, 2022. 185(10): p. 1777–1792. e21.

21. Biancalani, T., et al., Deep learning and alignment of spatially resolved single-cell transcriptomes with Tangram. Nature methods, 2021. 18(11): p. 1352–1362.

22. Ipsen, M. and A.S. Mikhailov, Evolutionary reconstruction of networks. Physical Review E, 2002. 66(4): p. 046109.

23. Van de Sande, B., et al., A scalable SCENIC workflow for single-cell gene regulatory network analysis. Nature protocols, 2020. 15(7): p. 2247–2276.

24. Ashuach, T., et al., MultiVI: deep generative model for the integration of multimodal data. Nature Methods, 2023. 20(8): p. 1222–1231.

25. Yan, F., et al., Single-cell multiomics decodes regulatory programs for mouse secondary palate development. Nature communications, 2024. 15(1): p. 821.

26. Setty, M., et al., Characterization of cell fate probabilities in single-cell data with Palantir. Nature biotechnology, 2019. 37(4): p. 451–460.

27. Ho, J., A. Jain, and P. Abbeel, Denoising diffusion probabilistic models. Advances in neural information processing systems, 2020. 33: p. 6840–6851.

28. Rombach, R., et al. High-resolution image synthesis with latent diffusion models. in Proceedings of the IEEE/CVF conference on computer vision and pattern recognition. 2022.

29. Pascanu, R., T. Mikolov, and Y. Bengio. On the difficulty of training recurrent neural networks. in International conference on machine learning. 2013. Pmlr.

30. Cui, H., et al., scGPT: toward building a foundation model for single-cell multi-omics using generative AI. Nature methods, 2024. 21(8): p. 1470–1480.

31. Wolf, F.A., P. Angerer, and F.J. Theis, SCANPY: large-scale single-cell gene expression data analysis. Genome biology, 2018. 19(1): p. 15.

32. Wolock, S.L., R. Lopez, and A.M. Klein, Scrublet: computational identification of cell doublets in single-cell transcriptomic data. Cell systems, 2019. 8(4): p. 281–291. e9.

33. Moerman, T., et al., GRNBoost2 and Arboreto: efficient and scalable inference of gene regulatory networks. Bioinformatics, 2019. 35(12): p. 2159–2161.

34. Imrichová, H., et al., i-cisTarget 2015 update: generalized cis-regulatory enrichment analysis in human, mouse and fly. Nucleic acids research, 2015. 43(W1): p. W57–W64.

