## Supplemental File for "CellPace: A temporal diffusion-forcing framework for simulation, interpolation and forecasting of single-cell dynamics"

### Supplementary Notes

#### Supplementary Note 1:

We performed systematic hyperparameter optimization for the four scRNA-seq lineages in [1] and the multiome dataset [2] using a validation set (separate from the test set).

Given the computational cost of generative modelling, an exhaustive grid search over the hyperparameters was infeasible. To explore this space efficiently, we used a two-stage Bayesian optimization strategy with Tree-structured Parzen Estimators (TPE) implemented in Optuna.

- Stage A: broad latent-space search (200 trials): In the first stage, we focused on structural and training hyperparameters. To maximize throughput, we evaluated models in the low-dimensional VAE latent space rather than in decoded gene space. We optimized architecture (for example, network depth, number of attention heads, hidden dimensions), training dynamics (learning rate, gradient clipping) and diffusion-specific parameters (SNR clipping, noise schedules). We used a median-pruner algorithm to terminate unpromising trials early (warm-up: 50% of training), based on 1,000 generated cells per validation stage.
- Stage B: fine-tuning in gene space (200 trials): We used the top 20 configurations from Stage A to seed the TPE sampler in the second stage. Here, models were fully decoded to gene-expression space to assess biological fidelity. This hierarchical design substantially reduced computational cost compared with a single-stage optimization directly in gene space while still guiding the search towards high-performing regions of the parameter space.

After hyperparameter tuning, we retrained the model on the combined training and validation sets for 20,000-40,000 iterations (depending on dataset size).

### Supplementary Note 2: Baseline models

**scDiffusion** [3] is a conditional latent diffusion model for generating single-cell RNA-seq profiles with specified cellular attributes. It consists of three sequentially trained modules. First, a deterministic autoencoder compresses log-transformed, library-size–normalized expression profiles into a compact, L2-normalized latent space using fully connected encoder and decoder networks with batch normalization and PReLU activations, trained with mean squared error loss. Second, a denoising diffusion model is trained in this latent space using a residual MLP with symmetric encoder–decoder architecture, skip connections, layer normalization, SiLU activations and sinusoidal timestep embeddings under a linear noise schedule. Third, a noise-conditional classifier is trained on latents noised at 500 timesteps to enable classifier guidance. For generation, the model samples from Gaussian noise and iteratively denoises via the reverse diffusion process. Classifier guidance is applied during the early denoising steps by shifting the predicted mean in the direction of the gradient of the conditioned log-probability. Continuous interpolation between cell labels can be achieved by partially noising latents from a source population and denoising them under weighted guidance towards multiple target classes, with optional filtering by predicted labels. Final expression profiles are obtained by passing the resulting latents through the pretrained decoder.

**cfDiffusion** [4] is a two-stage conditional latent diffusion model that reuses the deterministic autoencoder backbone of scDiffusion but replaces the separate noise-conditioned classifier with classifier-free guidance. In the diffusion stage, it extends scDiffusion's residual MLP backbone by incorporating both timestep and label embeddings: cell-type conditions are one-hot encoded, passed through a small conditioning network and injected into each residual block via adaptive normalization that combines multiplicative label modulation with additive time conditioning, while preserving the symmetric encoder-decoder structure, skip connections, layer normalization and SiLU activations. During training, the model learns both conditional and unconditional noise predictions by randomly dropping cell-type labels with a fixed probability under a linear noise schedule and predicting the added noise at each step. For generation, samples are drawn from a Gaussian prior in latent space and iteratively denoised with DDPM or DDIM sampling; classifier-free guidance is applied at every step by combining conditional and unconditional predictions with a fixed guidance strength ($w = 0.5$) to steer samples towards the target cell type. An optional feature-caching mechanism reuses intermediate activations from selected network layers across consecutive denoising steps to accelerate inference, and the final gene expression profiles are decoded with the autoencoder.

**ESCFD** [5] is a three-stage latent diffusion framework that combines an autoencoder, a masked autoencoder and a diffusion Transformer to model conditional single-cell RNA-seq distributions. First, a deterministic autoencoder is trained on expression profiles that are normalized to a fixed library size and log-transformed, mapping cells into a low-dimensional latent space and reconstructing counts via mean squared error loss. The resulting latent vectors are reshaped into small pseudo-images, enabling a second stage in which a Vision Transformer-based masked autoencoder learns structure in this latent representation by randomly masking most patches and reconstructing the missing content in a self-supervised manner. In the third stage, a diffusion Transformer (scDiT) operates on the same reshaped latent images, predicting a velocity field under AdaLN-Zero conditioning and an auxiliary feature-matching loss that encourages its internal representations to align with those of the frozen masked autoencoder. Class labels are embedded and used to condition the diffusion Transformer, allowing class-conditional generation. At inference, ESCFD samples latent images from a Gaussian prior, integrates the learned velocity field with an ODE-based sampler under classifier-free guidance, flattens the denoised latents back to vectors and decodes them through the pretrained autoencoder decoder to obtain generated log-normalized expression profiles.

**scNODE** [6] is a conditional generative model that combines a variational autoencoder with neural ordinary differential equations (ODE) to model developmental trajectories in continuous time. A VAE encoder maps gene expression profiles to a low-dimensional latent space with parameterized mean and standard deviation, a neural ODE defines a drift function in this latent space via a multilayer feedforward network with ReLU activations and a VAE decoder reconstructs gene expression from latent states. Training proceeds in two stages: the VAE is first fitted with mean squared error reconstruction loss to learn a compressed latent representation, after which the encoder, decoder and ODE drift network are jointly optimized using a composite objective that combines Sinkhorn divergence (optimal transport) between predicted and observed cell distributions at each timepoint with a dynamic regularization term enforcing consistency between ODE-evolved latents and VAE-encoded latents. For generation, unlike diffusion models or VAEs that can generate from noise, scNODE performs conditional generation: it encodes real initial cells, integrates the learned ODE forward in time and decodes the resulting states at any targeted timepoint.

**scIMF** [7] extends trajectory-based single-cell models by incorporating explicit cell–cell interactions through a McKean–Vlasov stochastic differential equation in a fixed 50-dimensional PCA space. Unlike scNODE, where cells evolve independently, the McKean–Vlasov framework models population-level dynamics in which each cell's drift depends on the empirical distribution of the entire population. This drift is decomposed into an intercellular component implemented by a Transformer with masked self-attention and an intracellular component implemented by a multilayer perceptron capturing cell-autonomous gene regulation; a constant diagonal diffusion term ($\sigma = 0.1$) adds stochasticity. scIMF is trained in a single stage with a composite loss combining Sinkhorn optimal transport to match predicted and observed cell distributions at each timepoint with an energy regularizer penalising large drift magnitudes. For generation, gene expression is zscore normalized and projected into PCA space at the initial timepoint; the learned McKean–Vlasov SDE is then integrated forward with an Euler–Maruyama scheme to the desired developmental stages, and the resulting latents are mapped back via the inverse PCA transform. As with scNODE, scIMF uses observed cells at the first timepoint as initial conditions to generate predictions at subsequent stages.

**CFGen** [8] is a two-stage conditional generative model that combines a deterministic autoencoder with flow matching in a learned latent space to model stage-conditional single-cell RNA-seq distributions. In the first stage, a fully connected autoencoder takes raw count matrices and internally applies log-normalization to encode gene expression into a latent representation, then decodes to negative binomial distribution parameters that reconstruct the original raw counts, with per-gene dispersion terms capturing overdispersion. The encoder is deterministic (no latent sampling or KL regularization), and library-size variation is handled via explicit size-factor inputs rather than being absorbed into the latent code. In the second stage, CFGen learns a time- and condition-dependent velocity field in this latent space using flow matching. Given an encoded cell (data latent), a latent noise sample and a conditioning label (such as somite stage or cell type), CFGen samples intermediate points along nearly straight-line paths between noise and data and trains a residual MLP to predict the target velocity that transports noise towards the data point. Conditioning information is assembled from sinusoidal time embeddings, size-factor embeddings and categorical embeddings for the discrete condition, which are summed and added to the hidden activations in each residual block as an additive shift, while the residual MLP retains the same overall structure as in the single-modal backbone. The flow-matching network is trained with a mean squared error loss between predicted and target velocities, with the autoencoder frozen, yielding a conditional latent flow that maps a Gaussian prior to the empirical cell distribution for each condition. For generation, CFGen samples latent vectors from a standard Gaussian prior, selects a categorical condition and an associated size factor, and numerically integrates the learned velocity field from noise ($t = 0$) to data ($t = 1$) with an adaptive ODE solver to obtain generated latents at the terminal time. These latents are passed through the pretrained decoder to obtain mean expression parameters, from which generated counts are sampled via the negative binomial likelihood. CFGen can optionally employ classifier-free guidance by randomly dropping condition labels during training and, at sampling time, combining conditional and unconditional velocity evaluations with a fixed guidance weight ($w = 1.0$) to sharpen adherence to the target condition.

### Supplementary Figures

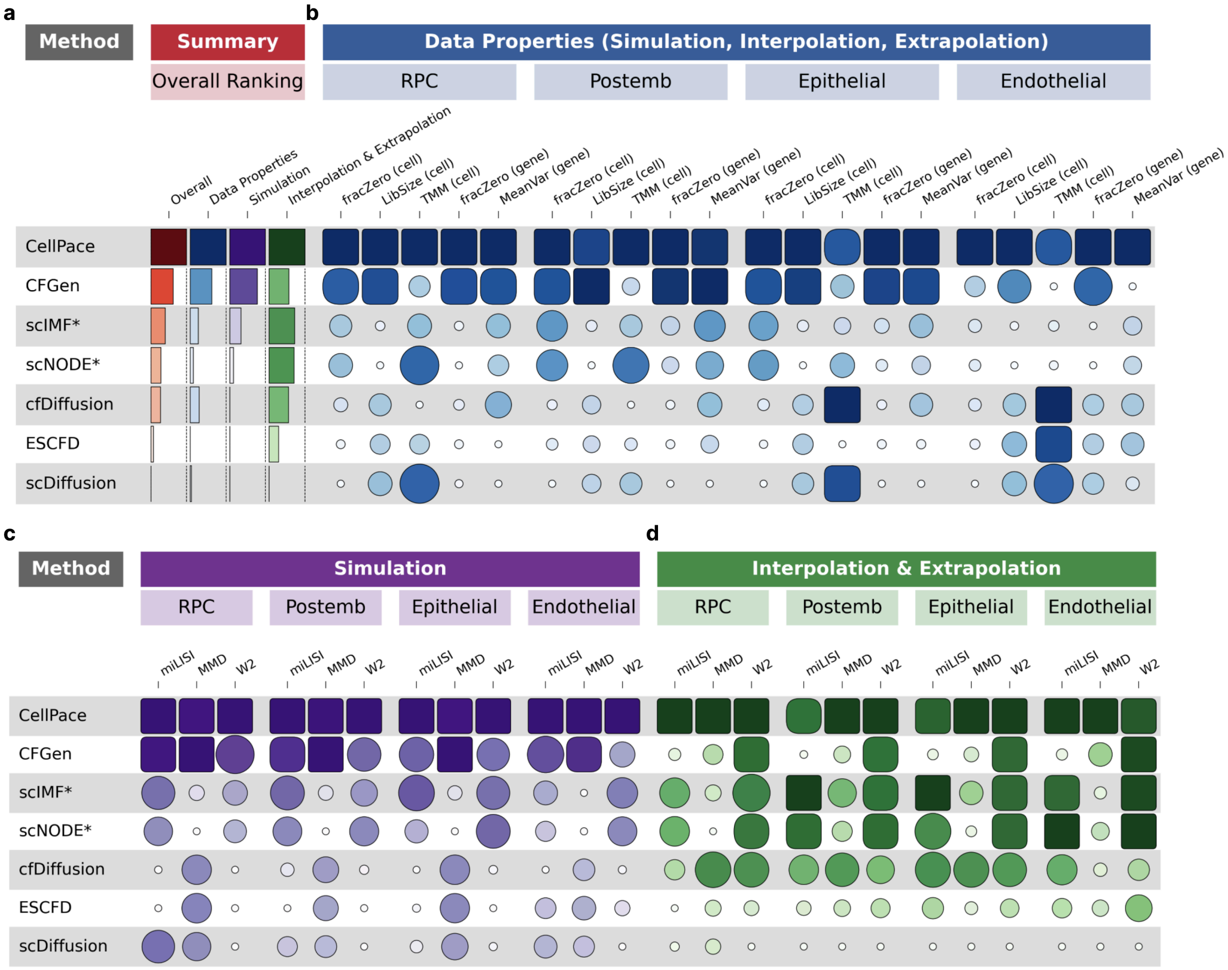

#### Supplementary Fig. 1 | Extended performance evaluation across metrics and datasets

Detailed performance evaluation corresponding to the summary ranking in Fig. 1f, visualized as a heatmap, where larger/darker shapes represent better performance (normalized scores 0-1). **a,** Summary scores aggregated for overall ranking across different methods. **b,** Data Properties (blue) assess the fidelity of statistical features. Metrics include cell-level properties (fraction of zeros per cell, library size per cell, and TMM normalization factors) and gene-level properties (fraction of zeros per gene and mean-variance relationships). **c,** Simulation (purple) evaluates generation quality on observed training stages using miLISI (measuring mixing in embedding space), MMD, and 2-Wasserstein distance in the PCA space. **d,** Interpolation & Extrapolation evaluates performance on held-out test stages using the same metrics.

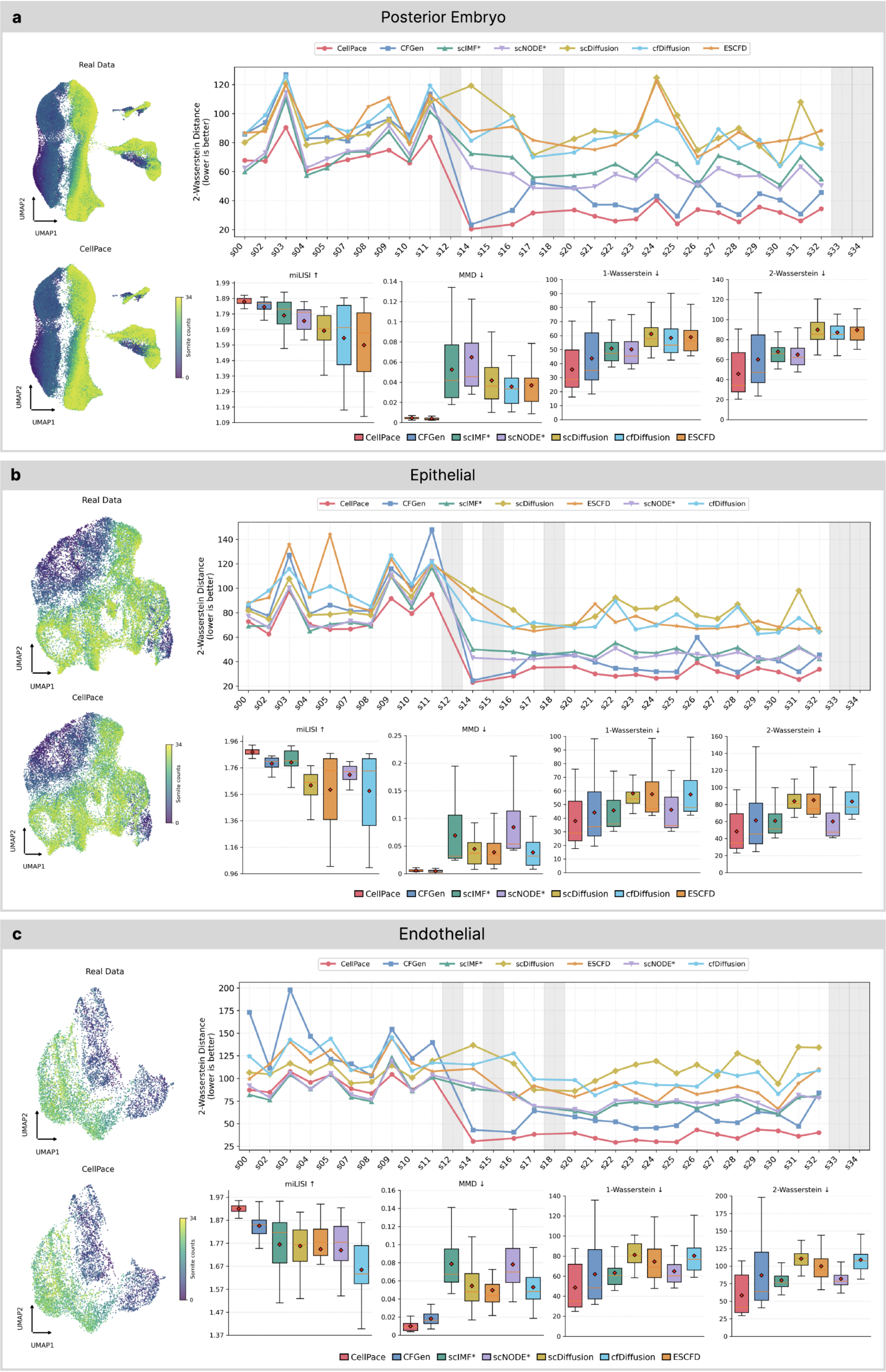
a
Posterior Embryo
Real Data
CellPace
UMAP1
UMAP2
Somite counts
CellPace CFGen scIMF* scNODE* scDiffusion cfDiffusion ESCFD
2-Wasserstein Distance (lower is better)
s00 s02 s03 s04 s05 s07 s08 s09 s10 s11 s12 s14 s15 s16 s17 s18 s20 s21 s22 s23 s24 s25 s26 s27 s28 s29 s30 s31 s32 s33 s34
miLISI ↑
MMD ↓
1-Wasserstein ↓
2-Wasserstein ↓
b
Epithelial
Real Data
CellPace
UMAP1
UMAP2
Somite counts
CellPace CFGen scIMF* scDiffusion ESCFD scNODE* cfDiffusion
2-Wasserstein Distance (lower is better)
miLISI ↑
MMD ↓
1-Wasserstein ↓
2-Wasserstein ↓
c
Endothelial
Real Data
CellPace
UMAP1
UMAP2
Somite counts
CellPace CFGen scIMF* scDiffusion ESCFD scNODE* cfDiffusion
2-Wasserstein Distance (lower is better)
miLISI ↑
MMD ↓
1-Wasserstein ↓
2-Wasserstein ↓

**Supplementary Fig. 2 | Extended evaluation of simulation across multiple lineages**

Evaluation of simulation quality for Posterior Embryo (**a**), Epithelial (**b**), and Endothelial (**c**) lineages. Left: Joint UMAP embedding of real and CellPace-generated cells (training stages only), colored by somite. Right (Top): Temporal consistency quantified by 2-Wasserstein distance across training somite stages. Right (Bottom): Box plots showing the distribution of performance metrics (miLISI, MMD, 1-Wasserstein, 2-Wasserstein) across all training stages. In all box plots, boxes denote the interquartile range (IQR), center lines mark medians, and whiskers extend to 1.5×IQR. Metrics evaluate generation quality on observed training data: higher miLISI indicates better mixing; lower MMD/Wasserstein distance indicate higher distributional fidelity.

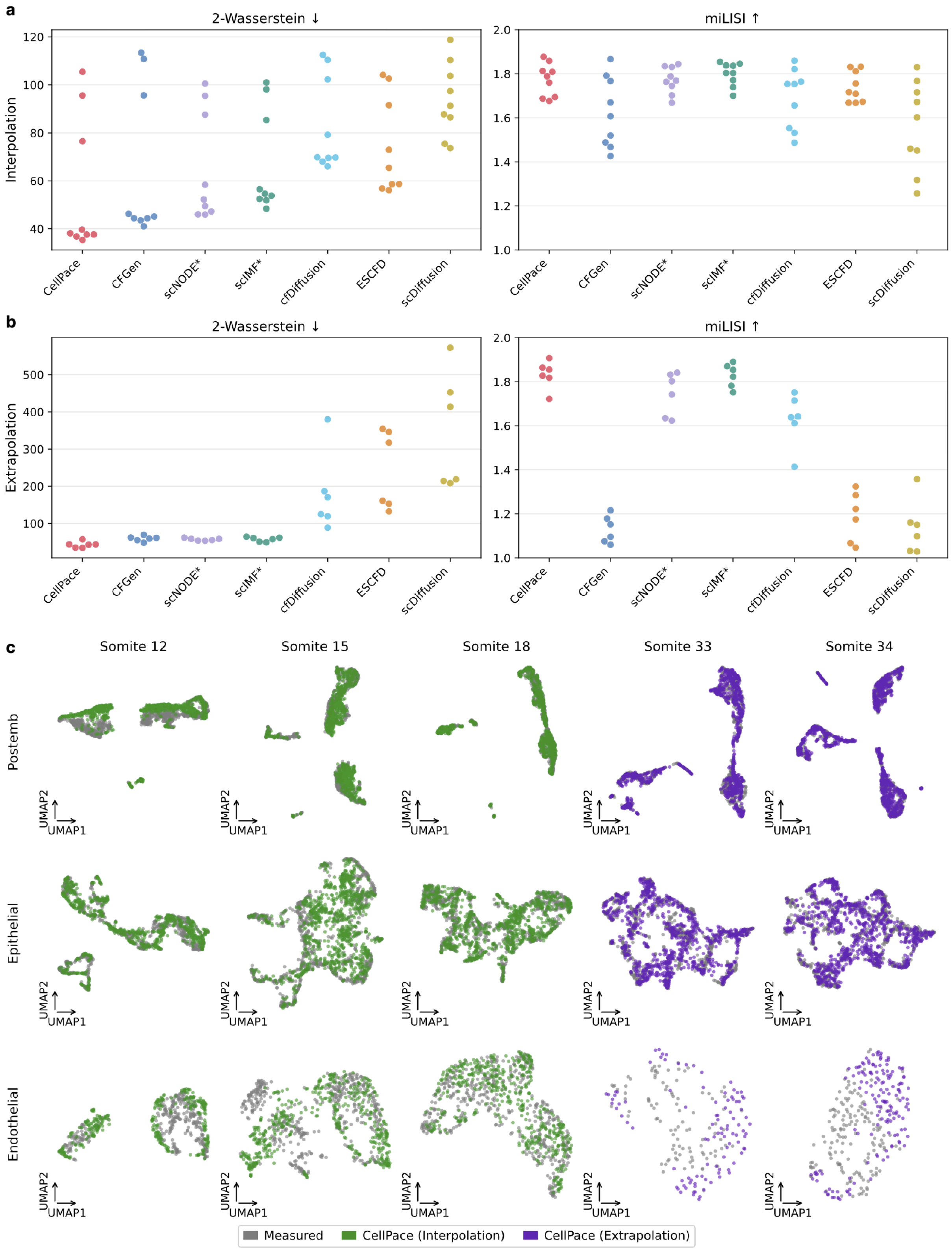

**Supplementary Fig. 3 | Extended evaluation of interpolation and extrapolation across multiple lineages**

**a,** Quantitative performance of different models in predicting the intermediate somite stages (Somite stages 12, 15, and 18). Swarm plots show the distribution of 2-Wasserstein distance and miLISI scores across Posterior Embryo, Epithelial, and Endothelial datasets, with each point representing one test stage from one dataset. **b,** Quantitative performance of different models in forecasting the future somite stages (Somite stages 33 and 34). Plots follow the same layout as in a, summarizing performance across the same three datasets. **c,** Joint UMAP visualizations for three datasets. Real (experimentally measured) cells (grey) are shown together with CellPace-generated cells for interpolation (green) and extrapolation (purple) across all five test somite stages in three datasets.

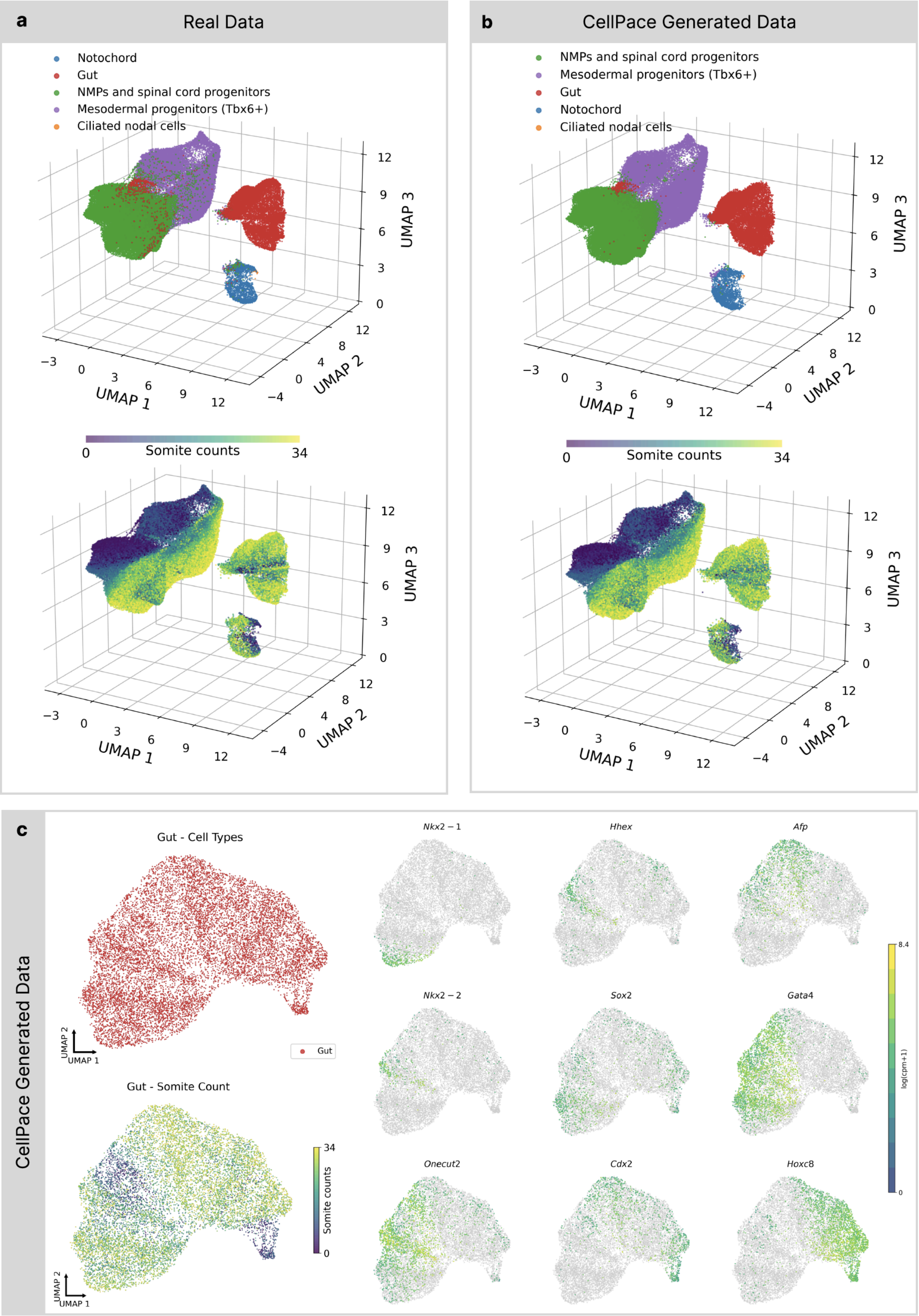
a
Real Data
Notochord
Gut
NMPs and spinal cord progenitors
Mesodermal progenitors (Tbx6+)
Ciliated nodal cells
UMAP 1
UMAP 2
UMAP 3
Somite counts
0
34
b
CellPace Generated Data
NMPs and spinal cord progenitors
Mesodermal progenitors (Tbx6+)
Gut
Notochord
Ciliated nodal cells
UMAP 1
UMAP 2
UMAP 3
Somite counts
0
34
c
CellPace Generated Data
Gut - Cell Types
Gut
Gut - Somite Count
Somite counts
UMAP 1
UMAP 2
Nkx2 − 1
Hhex
Afp
Nkx2 − 2
Sox2
Gata4
Onecut2
Cdx2
Hoxc8
log(cpm+1)
8.4
0

**Supplementary Fig. 4 | Extended analysis of manifold structure and lineage-specific transcriptional patterns in CellPace-generated posterior embryo samples**

**a,** 3D UMAP embedding of experimentally measured (real) data, colored by major lineages (Top) including notochord, gut, mesodermal progenitors, and neuromesodermal progenitors (NMPs), and colored by somite count (Bottom). **b,** CellPace-generated 3D UMAP embedding of CellPace-generated, shown using the same layout and color schemes as in **a**. **c,** Reconstruction of gut organogenesis. Left: Re-embedded 2D UMAP of CellPace-generated cells corresponding to the gut lineage. Right: Feature plots displaying log-normalized expression of marker genes defining foregut (*Nkx2-1, Hhex, Nkx2-2, Sox2*), midgut/hindgut (*Cdx2, Hoxc8*), and liver/pancreas progenitors (*Afp, Gata4, Onecut2*) visualized on the re-computed gut manifold.

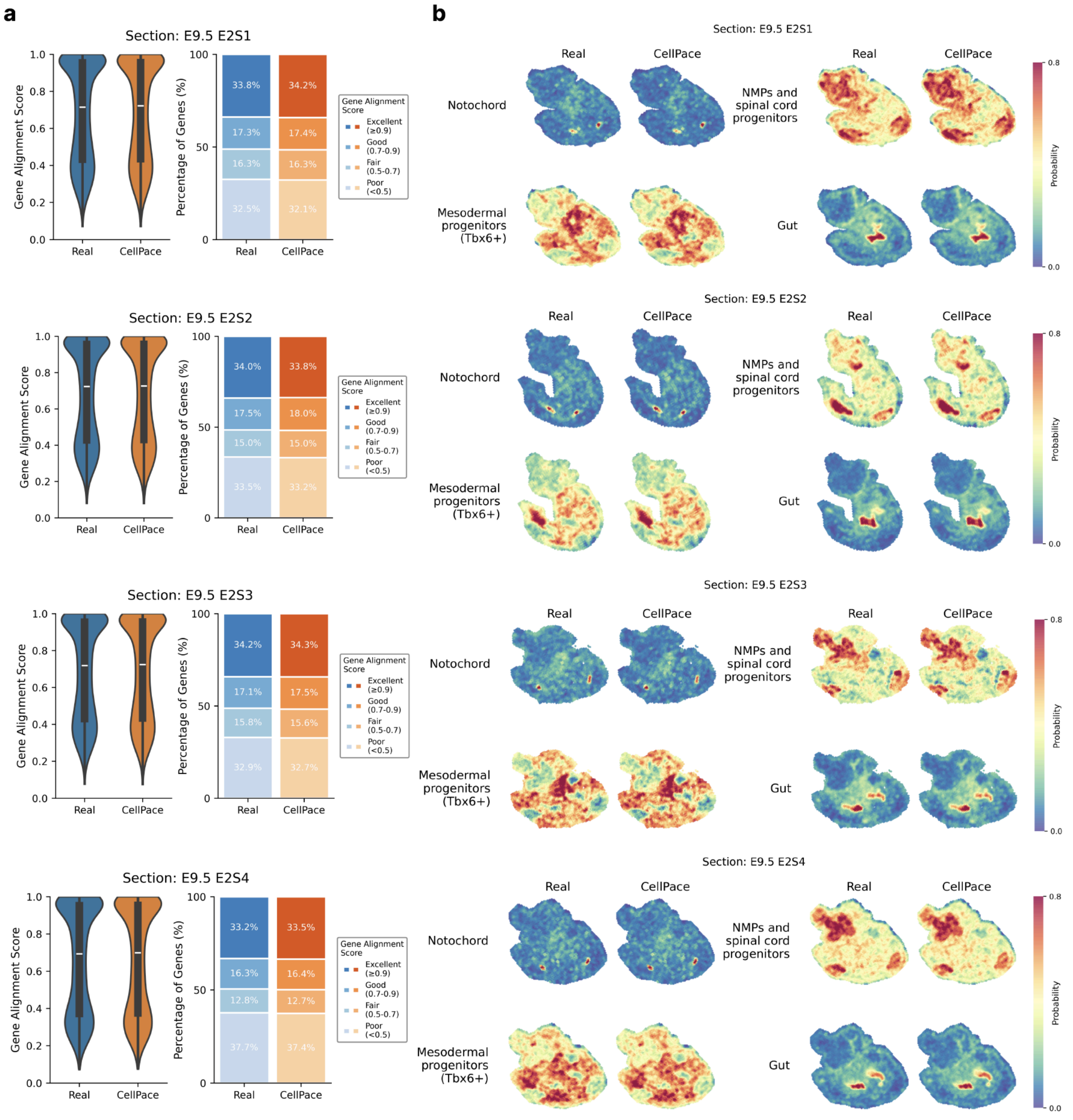

**Supplementary Fig. 5 | Extended spatial mapping analysis of CellPace-generated posterior embryo samples across multiple MOSTA sections**

**a,** Extended spatial mapping quality assessment. Evaluation of single-cell mapping to four additional MOSTA sections (E9.5 E2S1, E2S2, E2S3, and E2S4) using Tangram. For each section, violin plots (left) show the distribution of gene alignment scores for real and CellPace-generated data, while stacked bar charts (right) summarize the proportion of genes across alignment score categories (Excellent ≥ 0.9, Good 0.7–0.9, Fair 0.5–0.7, and Poor < 0.5). **b,** Spatial localization across multiple sections. Spatial probability maps for representative lineages (notochord, neuromesodermal progenitors, mesodermal progenitors, and gut) reconstructed on sections E2S1, E2S2, E2S3, and E2S4, shown for real and CellPace-generated data.

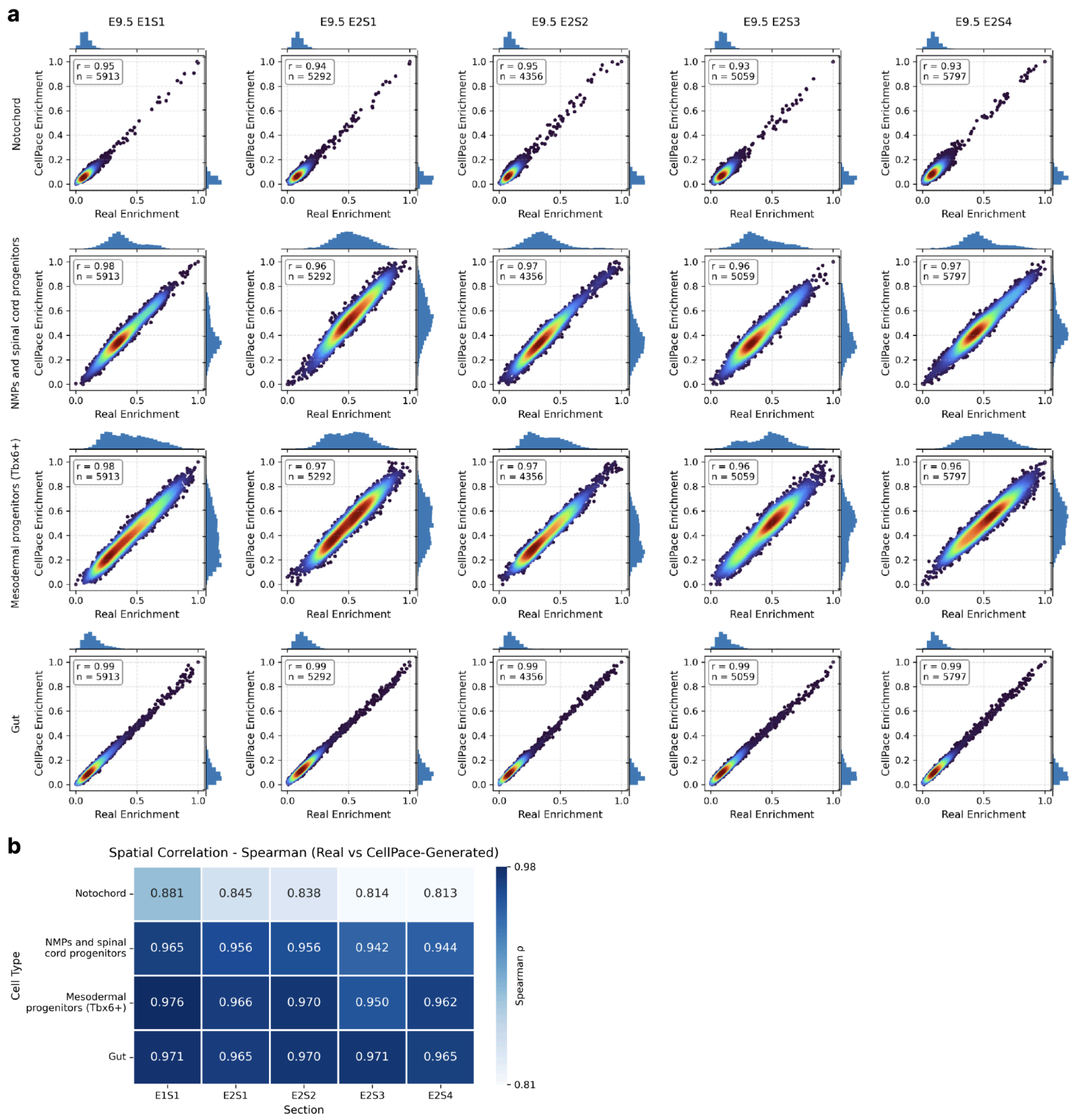

**Supplementary Fig. 6 | Extended quantitative comparison of spatial enrichment scores derived from real and CellPace-generated data**

**a,** Quantitative comparison of spatial enrichment scores. Density-colored scatter plots with marginal histograms comparing relative spatial enrichment scores derived from real (x-axis) and CellPace-generated (y-axis) data across four cell types and five anatomical sections. Enrichment scores are computed from Tangram cell-type mapping outputs by k-nearest neighbor smoothing in physical space followed by min-max normalization to [0, 1]. Each point represents one spatial spot ($n \approx 4300 - 5900$ spots per section), and Pearson correlation coefficients (r) are reported for each cell type–section combination. **b,** Spatial rank correlation of enrichment scores. Heatmap showing Spearman correlation coefficients ($\rho$) between real and CellPace-derived enrichment score vectors for each cell type across sections, computed over spatial spots within each section.

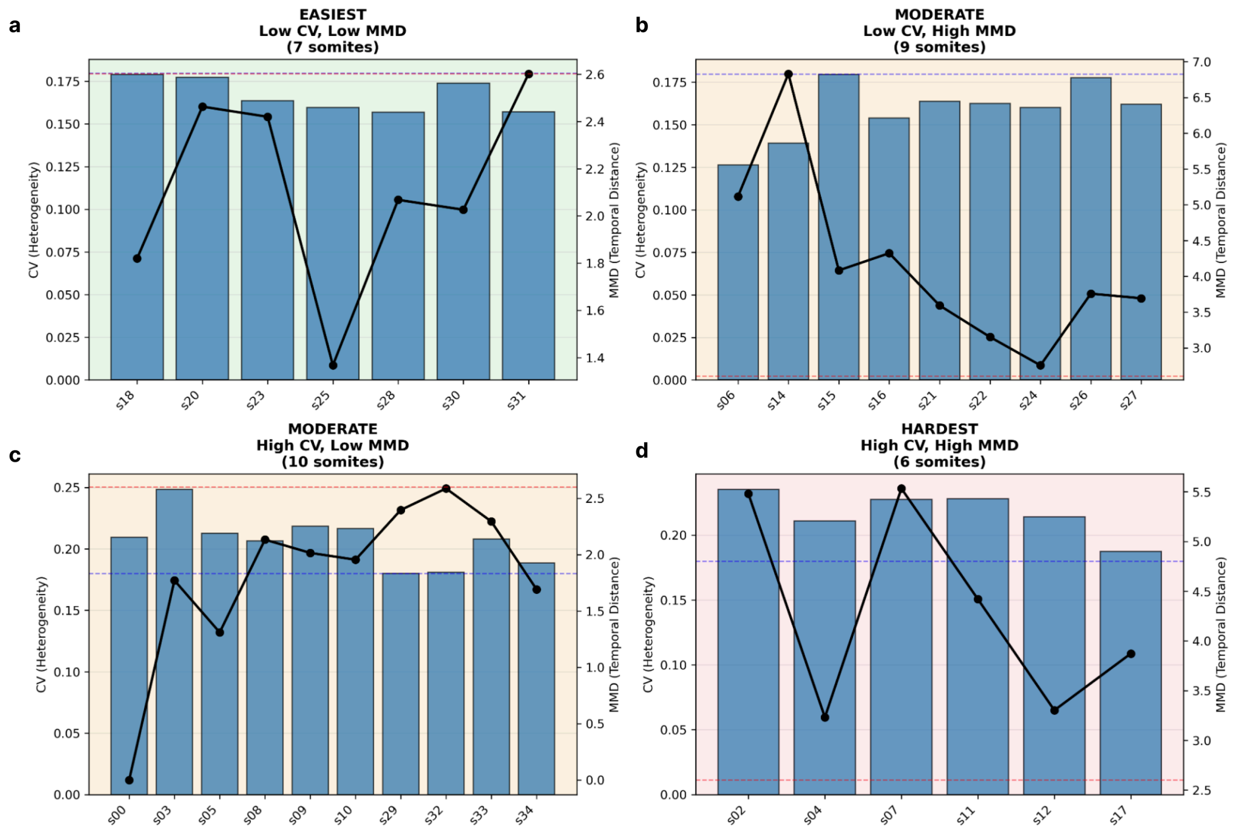

**Supplementary Fig. 7 | Data-driven stratification of somite difficulty based on intrinsic heterogeneity and temporal dynamics**

Somite stages were analyzed using the top 3,000 highly variable genes (HVGs) in PCA space (top 50 components) and grouped into four difficulty categories based on two intrinsic metrics. CV (Coefficient of Variation, blue bars, left y-axis) quantifies intra-somite cellular heterogeneity (normalized standard deviation of pairwise distances). MMD (Maximum Mean Discrepancy, black line, right y-axis) measures the magnitude of the developmental transition (Euclidean distance between centroids of consecutive somite stages). Blue and red dashed lines indicate median thresholds (blue: CV=0.180; red: MMD=2.60). The resulting categories are: EASIEST (Low CV/Low MMD; green, n=7) representing stable populations with small transitions; MODERATE (orange, n=19 total) comprising Low CV/High MMD (n=9) and High CV/Low MMD (n=10), representing intermediate difficulty; and HARDEST (High CV/High MMD; red, n=6) representing highly variable populations with large developmental jumps. Based on this stratification, test somite stages were selected to span the full difficulty spectrum: Somite 18 (Easiest), Somite 15 (Moderate), and Somite 12 (Hardest) for interpolation. For extrapolation, Somite stages 33 and 34 fall into the Moderate category (High CV/Low MMD), representing highly heterogeneous populations with small transitions between consecutive stages.

### Supplementary Tables

#### Supplementary Table 1 | Distributional similarity metrics on training developmental stages.

| dataset | Endothelial | | | Epithelial | | | Postemb | | | RPC | | |
|---|---|---|---|---|---|---|---|---|---|---|---|---|
| metric | MMD | W2 | miLISI | MMD | W2 | miLISI | MMD | W2 | miLISI | MMD | W2 | miLISI |
| **CFGen [8]** | 0.02 | 105.27 | 1.85 | 0.00 | 78.69 | 1.79 | 0.00 | 76.36 | 1.84 | 0.00 | 64.68 | 1.88 |
| **CellPace** | 0.01 | 86.60 | 1.91 | 0.01 | 66.49 | 1.88 | 0.00 | 64.09 | 1.87 | 0.01 | 57.44 | 1.89 |
| **ESCFD [5]** | 0.05 | 110.95 | 1.74 | 0.04 | 99.30 | 1.59 | 0.04 | 105.22 | 1.59 | 0.04 | 89.92 | 1.46 |
| **cfDiffusion [4]** | 0.05 | 121.37 | 1.65 | 0.04 | 97.86 | 1.58 | 0.04 | 104.39 | 1.63 | 0.04 | 89.63 | 1.46 |
| **scDiffusion [3]** | 0.05 | 122.86 | 1.76 | 0.04 | 97.57 | 1.62 | 0.04 | 106.64 | 1.68 | 0.04 | 88.69 | 1.74 |
| **scIMF*[7]** | 0.08 | 86.77 | 1.76 | 0.07 | 69.22 | 1.80 | 0.05 | 75.56 | 1.78 | 0.07 | 67.48 | 1.75 |
| **scNODE* [6]** | 0.08 | 88.28 | 1.74 | 0.08 | 68.07 | 1.70 | 0.06 | 72.54 | 1.74 | 0.09 | 68.70 | 1.70 |

Note: The table reports distributional similarity metrics evaluated on training developmental stages, including miLISI (median Local Inverse Simpson's Index computed on a joint t-SNE embedding; higher is better), MMD (data) (Maximum Mean Discrepancy computed in expression space; lower is better), and W2 (2-Wasserstein distance computed in PCA space; lower is better). Asterisk (*) denotes methods that require real cells from the initial timepoint as input, rather than generating from noise. This table corresponds to Supplementary Figure 1c.

**Supplementary Table 2 | Distributional similarity metrics on held-out test developmental stages**

| dataset | Endothelial | | | Epithelial | | | Postemb | | | RPC | | |
|---|---|---|---|---|---|---|---|---|---|---|---|---|
| **metric** | **MMD** | **W2** | **miLISI** | **MMD** | **W2** | **miLISI** | **MMD** | **W2** | **miLISI** | **MMD** | **W2** | **miLISI** |
| **CFGen** | 0.07 | 82.52 | 1.48 | 0.06 | 76.23 | 1.37 | 0.06 | 76.56 | 1.42 | 0.06 | 69.94 | 1.44 |
| **CellPace** | 0.02 | 75.15 | 1.81 | 0.01 | 70.50 | 1.81 | 0.01 | 64.91 | 1.78 | 0.01 | 63.16 | 1.80 |
| **ESCFD** | 0.08 | 149.64 | 1.55 | 0.07 | 169.78 | 1.51 | 0.06 | 156.52 | 1.50 | 0.07 | 164.04 | 1.39 |
| **cfDiffusion** | 0.09 | 174.13 | 1.64 | 0.03 | 113.36 | 1.69 | 0.03 | 128.00 | 1.66 | 0.03 | 97.21 | 1.53 |
| **scDiffusion** | 0.10 | 227.44 | 1.44 | 0.08 | 209.45 | 1.32 | 0.07 | 208.13 | 1.41 | 0.07 | 186.56 | 1.41 |
| **scIMF*** | 0.09 | 73.63 | 1.76 | 0.05 | 70.40 | 1.83 | 0.04 | 71.92 | 1.84 | 0.07 | 79.52 | 1.63 |
| **scNODE*** | 0.08 | 72.09 | 1.79 | 0.07 | 69.64 | 1.70 | 0.05 | 68.86 | 1.80 | 0.09 | 75.28 | 1.63 |

Note: The table reports distributional similarity metrics evaluated on test developmental stages, same metrics as Supplementary Table 1. Asterisk (*) denotes methods that require real cells from the initial timepoint as input, rather than generating from noise. This table corresponds to Supplementary Figure 1d.

**Supplementary Table 3 | Temporal correlation of regulon activity dynamics between real and CellPace-generated cells across developmental stages**

| Regulon | Pearson_r | p_value | Num of somite stages |
|---|---|---|---|
| Etv4(+) | 0.988751 | 1.75E-25 | 31 |
| Hoxc11(+) | 0.986622 | 2.13E-24 | 31 |
| Hoxd11(+) | 0.974204 | 2.68E-20 | 31 |
| Msx1(+) | 0.97009 | 2.23E-19 | 31 |
| Hoxd10(+) | 0.967802 | 6.4E-19 | 31 |
| Hoxa10(+) | 0.964906 | 2.19E-18 | 31 |
| Hoxa6(+) | 0.955369 | 6.73E-17 | 31 |
| Bcl11a(+) | 0.952782 | 1.5E-16 | 31 |
| Hoxc10(+) | 0.952408 | 1.68E-16 | 31 |
| Hoxc9(+) | 0.951027 | 2.52E-16 | 31 |
| Hoxc5(+) | 0.946097 | 9.79E-16 | 31 |
| Nkx1-2(+) | 0.941989 | 2.76E-15 | 31 |
| Hoxa11(+) | 0.932448 | 2.36E-14 | 31 |
| Hoxc8(+) | 0.932281 | 2.45E-14 | 31 |
| Tcf7l1(+) | 0.92131 | 2.01E-13 | 31 |
| Pou3f2(+) | 0.919238 | 2.89E-13 | 31 |
| Elk3(+) | 0.919081 | 2.97E-13 | 31 |
| Hoxa9(+) | 0.90604 | 2.38E-12 | 31 |
| Hoxd9(+) | 0.9049 | 2.81E-12 | 31 |
| Hoxa7(+) | 0.889982 | 2.11E-11 | 31 |
| Nkx2-1(+) | 0.889903 | 2.13E-11 | 31 |
| Pax3(+) | 0.881054 | 6.16E-11 | 31 |
| Sox9(+) | 0.876708 | 1.01E-10 | 31 |
| Cdx2(+) | 0.874376 | 1.3E-10 | 31 |
| Rarb(+) | 0.871259 | 1.82E-10 | 31 |
| Hivep2(+) | 0.865171 | 3.41E-10 | 31 |
| Sox10(+) | 0.853265 | 1.07E-09 | 31 |
| Sox2(+) | 0.850862 | 1.34E-09 | 31 |
| Ets2(+) | 0.849349 | 1.53E-09 | 31 |
| Pax9(+) | 0.830933 | 7.19E-09 | 31 |
| Foxa3(+) | 0.827376 | 9.5E-09 | 31 |
| Gli2(+) | 0.823747 | 1.25E-08 | 31 |
| Hoxb1(+) | 0.815771 | 2.25E-08 | 31 |
| Meis1(+) | 0.810773 | 3.21E-08 | 31 |
| Ets1(+) | 0.810594 | 3.25E-08 | 31 |
| Sox4(+) | 0.803914 | 5.13E-08 | 31 |
| Lef1(+) | 0.799283 | 6.97E-08 | 31 |
| Foxc1(+) | 0.798466 | 7.35E-08 | 31 |

| | | | |
|---|---|---|---|
| Nr3c1(+) | 0.793509 | 1.01E-07 | 31 |
| Foxa1(+) | 0.781105 | 2.16E-07 | 31 |
| Tbx6(+) | 0.775546 | 2.99E-07 | 31 |
| Meox1(+) | 0.764183 | 5.64E-07 | 31 |
| Nkx6-2(+) | 0.761493 | 6.53E-07 | 31 |
| Tead4(+) | 0.741675 | 1.8E-06 | 31 |
| Gata3(+) | 0.740455 | 1.92E-06 | 31 |
| Runx1(+) | 0.737944 | 2.16E-06 | 31 |
| Gata4(+) | 0.726925 | 3.63E-06 | 31 |
| Pdx1(+) | 0.724702 | 4.02E-06 | 31 |
| Onecut2(+) | 0.703128 | 1.03E-05 | 31 |
| Gata6(+) | 0.69077 | 1.7E-05 | 31 |
| Pou6f1(+) | 0.690317 | 1.73E-05 | 31 |
| Hnf4a(+) | 0.68948 | 1.79E-05 | 31 |
| Otx1(+) | 0.68356 | 2.25E-05 | 31 |
| Bach2(+) | 0.680732 | 2.51E-05 | 31 |
| Dbx2(+) | 0.632809 | 0.000133 | 31 |
| Hnf1b(+) | 0.615157 | 0.000231 | 31 |
| Ikzf2(+) | 0.607677 | 0.000288 | 31 |
| Gata5(+) | 0.605799 | 0.000304 | 31 |
| Pitx1(+) | 0.535498 | 0.001907 | 31 |
| Lmx1b(+) | 0.526375 | 0.002352 | 31 |
| Hnf4g(+) | 0.522568 | 0.002563 | 31 |
| Nkx6-1(+) | 0.416806 | 0.019669 | 31 |
| Foxa2(+) | 0.369857 | 0.040568 | 31 |
| Foxo1(+) | 0.335544 | 0.064984 | 31 |
| Elf1(+) | 0.294395 | 0.107919 | 31 |
| Sox17(+) | 0.243143 | 0.1875 | 31 |
| Nfia(+) | 0.208551 | 0.260216 | 31 |
| Foxc2(+) | -0.36706 | 0.042232 | 31 |

Note: Gene regulatory networks were inferred independently from real and CellPace-generated cells using the pySCENIC pipeline [9], consisting of GRNBoost2 for transcription factor–target co-expression inference, cisTarget for motif-based pruning of indirect targets, and AUCell for per-cell regulon activity scoring. For each regulon present in both networks, mean regulon activity was computed per somite stage, and Pearson correlation coefficients were calculated between the real and CellPace temporal activity profiles across all shared stages. Table columns: Regulon, transcription factor defining the regulon; Pearson_r, correlation coefficient between real and CellPace mean activity trajectories; p_value, statistical significance of the correlation; and number of somite stages used in the analysis. Rows are sorted by Pearson_r in descending order. Analysis includes both training and held-out test stages.

### Reference

1. Qiu, C., et al., *A single-cell time-lapse of mouse prenatal development from gastrula to birth.* Nature, 2024. **626**(8001): p. 1084-1093.
2. Yan, F., et al., *Single-cell multiomics decodes regulatory programs for mouse secondary palate development.* Nature communications, 2024. **15**(1): p. 821.
3. Luo, E., et al., *scDiffusion: conditional generation of high-quality single-cell data using diffusion model.* Bioinformatics, 2024. **40**(9): p. btae518.
4. Zhang, T., et al., *cfDiffusion: diffusion-based efficient generation of high quality scRNA-seq data with classifier-free guidance.* Briefings in Bioinformatics, 2025. **26**(1): p. bbaf071.
5. Li, S., et al. *ESCFD: Probabilistic Flow Diffusion Model for Accelerated High-Quality Single-Cell RNA-seq Data Synthesis*. in *Proceedings of the 31st ACM SIGKDD Conference on Knowledge Discovery and Data Mining V. 2*. 2025.
6. Zhang, J., et al., *scNODE: generative model for temporal single cell transcriptomic data prediction.* Bioinformatics, 2024. **40**(Supplement_2): p. ii146-ii154.
7. Jiang, Q., et al., *Learning collective multi-cellular dynamics from temporal scRNA-seq via a transformer-enhanced Neural SDE.* arXiv preprint arXiv:2505.16492, 2025.
8. Palma, A., et al., *Generating multi-modal and multi-attribute single-cell counts with CFGen.* arXiv preprint arXiv:2407.11734, 2024.
9. Van de Sande, B., et al., *A scalable SCENIC workflow for single-cell gene regulatory network analysis.* Nature protocols, 2020. **15**(7): p. 2247-2276.
